# Viruses, Proviruses and Satellites from Asgard Archaea Enrichments Reveal Complex Microbial Interactions

**DOI:** 10.64898/2026.08.12.739948

**Authors:** Julia Meltzer, Xabier Vázquez-Campos, Matthew D. Johnson, Thomas Litfin, Veronika Valova, Daniel Luque, Maria-Calliope Syrmalis, Keiran Rowell, Liam Hewitt, Bindusmita Paul, Katharine A. Michie, Miranda E. Pitt, Debnath Ghosal, Belinda C. Ferrari, Brendan P. Burns

## Abstract

Asgard archaea are the closest known relatives of eukaryotes and are central to models of eukaryogenesis involving archaeal–bacterial symbiosis, yet the contribution of viruses remains unexplored. Here, for the first time, we visualised unique viruses associated with Asgard archaeal cells. Additionally, we identified the first putative Asgard-archaeal virus satellite, which exhibited genomic interactions involving a bacterium, *Stromatodesulfovibrio nilemahensis*, providing evidence of a virus-mediated interaction between an Asgard archaeon and a bacterium. Additionally, novel proviruses of *S. nilemahensis* displayed distinct genomic features where predictions of alternate recombination sites suggested the acquisition of horizontally acquired genes associated with biofilm formation. Further, we comprehensively characterise (pro)viruses associated with this co-culture using high resolution cryo-electron tomography, proximity ligation (Hi-C), and metagenomics. Together, these findings expand the known diversity of Asgard archaeal viruses and establish a foundation for investigating the role of viruses in microbial symbiosis relevant to the emergence of eukaryotic life.

## Main

Asgard archaea have been widely regarded as the closest known evolutionary ancestors to eukaryotes. Metagenome-assembled genomes (MAGs) from Asgard lineages encode for a high number of eukaryotic signature proteins (ESPs)^1–4^, and phylogenomic reconstructions of the Tree of Life place eukaryotes as emerging from within the Asgard superphylum^3–5^. Despite the central importance of ancient Asgard archaea in the evolution and emergence of eukaryotes (termed eukaryogenesis), the cellular and ecological processes contributing to eukaryogenesis remains unknown^6^.

Leading models of eukaryogenesis propose that interactions between ancestral Asgard archaea and bacterial partners were critical to the development of early eukaryotes^1–3,6^. Specifically, these theories propose a symbiotic relationship between an ancient Asgard archaeon and an alphaproteobacterium, with many recent hypotheses including a fermentative deltaproteobacteria symbiont in a tripartite association^7,8^. Horizontal gene transfer (HGT) is implicated as a playing a major role in these endosymbiotic interactions, having contributed to the chimeric evolutionary origins of the eukaryotic genome^6,9,10^. Mobile genetic elements, including viruses, may have facilitated these symbiotic interactions and gene transfer events, as supported by evidence for cross-domain gene transfer associated with Asgard mobile genetic elements^11^. However, the role of viruses infecting Asgard archaea and their potential syntrophic partners is yet to be revealed, with the Asgard archaeal mobilome underexplored.

To date, the few viruses associated with Asgard archaea have been identified from metagenomic analyses^12–15^. While these studies revealed substantial viral diversity, they provided limited insights into viral structure, infection dynamics or host-interactions. Acquiring such new knowledge requires complementary, cultivation-based approaches. Stable, high abundance enrichment cultures of the Asgard archaea phylum (recently classified as *Promethearchaeota*^16^) have only recently been established, primarily from *Lokiarchaeia* lineages^17–19^. These cultivated Asgard archaeal representatives display complex cellular architecture and cell biology. In addition, they exhibit potential syntrophic interactions with bacterial partners, specifically sulfate reducing bacteria, providing experimental systems in which archaeal-bacterial interactions can be investigated directly. Despite these advances, viruses have not been characterised from these Asgard archaeal enrichments, leaving a major gap in our understanding of the Asgard mobilome.

We recently established enrichment cultures seeded with living microbial mats from Shark Bay (*Gathaagudu*), Australia, containing a high abundance of the novel *Lokiarchaeia* representative, *Nerearchaeum marumarumayae*, co-cultured with the sulfate-reducing bacterium *Stromatodesulfovibrio nilemahensis*^20^. Shark Bay microbial mats are well established analogues of Proterozoic ecosystems^21^, an era that coincides with the proposed timing of eukaryogenesis. Ancient microbial mats have also been proposed as a potential niche for eukaryogenesis^22^. Asgard archaea and sulfate-reducing bacteria are prominent in Shark Bay microbial mats^23,24^, where dense and diverse microbial and viral assemblages may facilitate cross-domain and viral interactions^25,26^. Therefore, stable enrichments of *N. marumarumayae* and *S. nilemahensis* originating from microbial mats provide a unique opportunity to examine host-virus interactions in detail.

Here, we present the first integrated characterisation of viruses infecting an Asgard archaeon in an enrichment culture. Using a combination of cryo-electron tomography (cryo-ET), proximity ligation (Hi-C) and metagenomics, we identify and visualise previously undescribed viruses associated with *N. marumarumayae,* and characterise four novel (pro)viruses associated with *S. nilemahensis.* We further uncover evidence for a putative Asgard archaea virus satellite and virus-mediated interactions between archaeal and bacterial members in co-culture. These findings expand the known diversity of the Asgard archaeal virome and provide insights into the potential role of viruses in the evolution of eukaryotes.

## Results

### Visualisation of novel Asgard archaeal viruses reveals diverse morphologies

While enrichment cultures of *N. marumarumayae* and *S. nilemahensis* have previously revealed evidence of metabolic syntrophy and direct cell-cell interactions^20^, their associated virome has not been characterised. To investigate the viral component of these enrichment cultures, virus-like particles (VLPs) were first examined by negative-stain transmission electron microscopy (TEM). A diverse assemblage of VLP morphologies was observed, including cosmopolitan archaeal and bacterial morphotypes (Fig. 1). These included head-tail structures characteristic of the class *Caudoviricetes*^27–29^, with both sheathed, contractile tails and sheathless, non-contractile variants (Fig. 1A). Spindle-shaped VLPs associated with archaea were also identified (Fig. 1B), as well as putative capsids of elongated or prolate morphotypes (Fig. 1C) and diverse decorated capsids (Fig. 1D). Some putative capsids had short tail-like structures, consistent with podovirus morphotypes within the *Caudoviricetes* class (Fig. 1E). Long filamentous structures were also identified, some of which may represent filamentous viruses (Fig. 1F). In addition to the virus-like particles described, numerous diverse unidentified extracellular particles were observed, potentially novel VLPs or extracellular entities (Fig. 1G).

**Figure 1.**
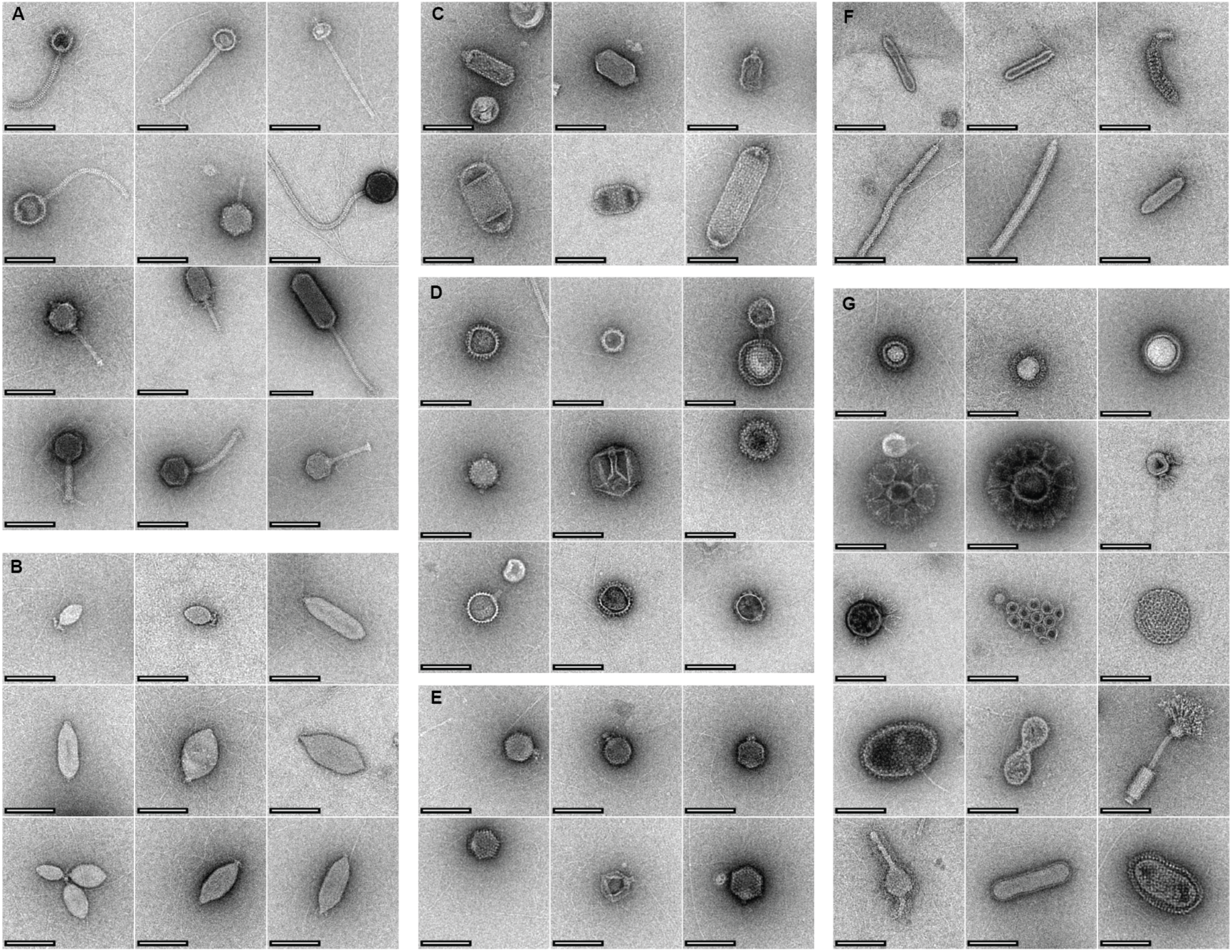
Diversity of virus-like particle morphotypes in Asgard archaeal enrichment cultures. Negative-stain transmission electron microscopy revealed diverse virus-like particles in low abundance Asgard archaeal enrichment cultures, including contractile and non-contractile head-tail phage morphotypes (A), spindle virus morphotypes (B), putative elongated or prolate capsids (C), and decorated capsids (D). Some putative capsids were associated with short tail-like structures, representative of podovirus morphotypes (E). Long filamentous structures were identified, some potentially representing filamentous viruses (F). Numerous diverse, unidentified extracellular particles were also detected (G). Scalebar = 100 nm.

Because the low biomass of the enrichment cultures limited particle recovery, additional cultures containing lower relative abundance of *N. marumarumayae* were included to obtain sufficient VLP concentrations for imaging. Consequently, the TEM observations provide an important overview of viral diversity of these enrichment cultures, however they did not allow for assignment of individual VLP morphotypes to a specific host.

Building on these observations we employed cryo-ET, a high-resolution imaging method capable of capturing features in near-native conditions (without fixing or staining). Cryo-ET imaging of the enriched cultures for viruses associated with *N. marumarumayae* or *S. nilemahensis*^20^ revealed that VLPs were associated with *N. marumarumayae* cell surfaces (Fig. 2 and Supplementary Video 1), whereas no virus particles were detected in association with *S. nilemahensis*. Two distinct VLP morphotypes were identified on *N. marumarumayae*: a sheathless head-tail phage resembling members of *Caudoviricetes*, and a spindle-shaped virus with morphology similar to members of the families *Fuselloviridae*^30^, *Thaspiviridae*^31^ and *Halspiviridae*^32^.

**Figure 2.**
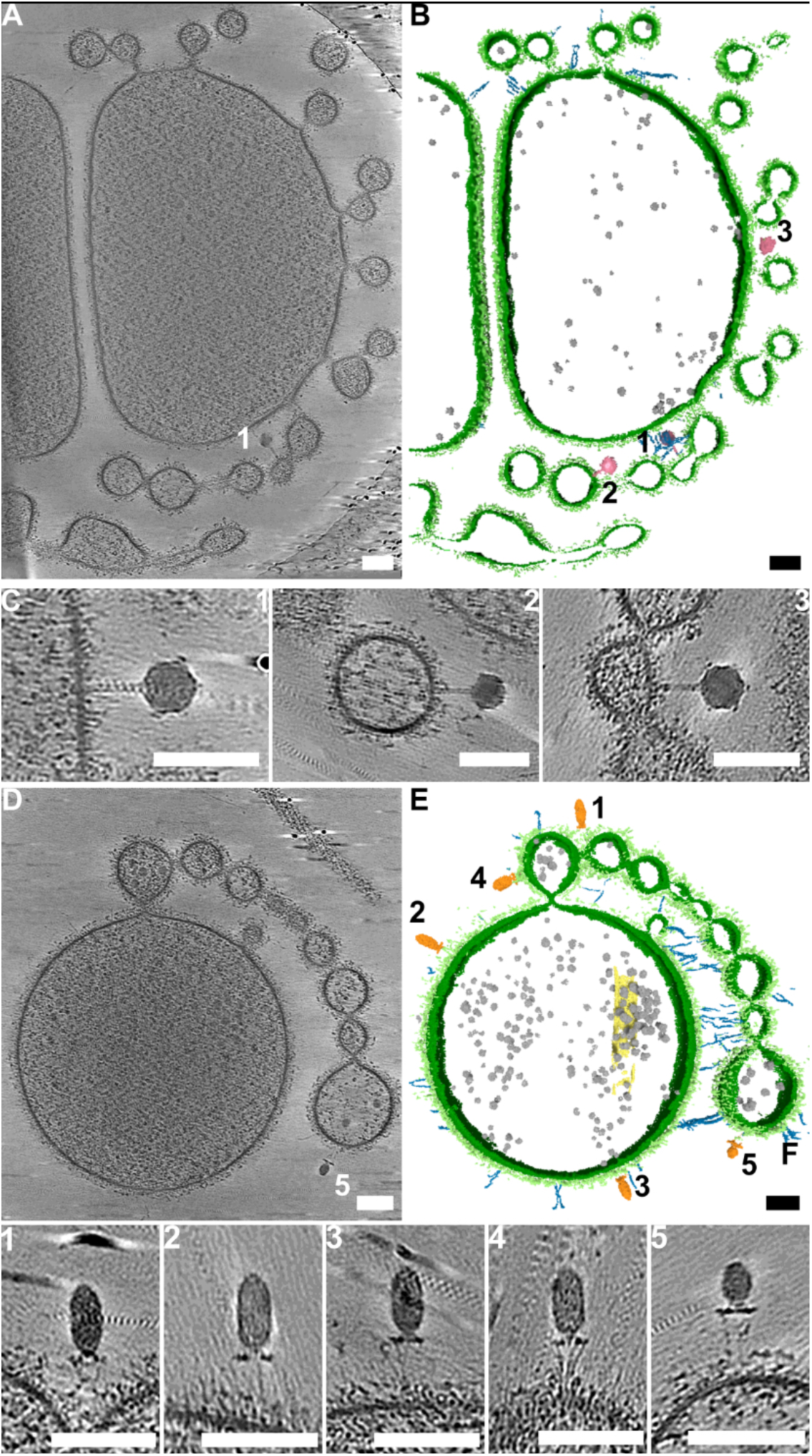
Cryo-ET images of VLPs associated with *Nerearchaeum marumarumayae* cells. **(A)** 2D-slice through at 3D tomogram of a *N. marumarumayae* cell (10-slices averaged). **(B)** Segmented volume of data shown in (A), showing inner membrane (dark green), outer surface layer (light green), ribosomes (grey), and interacting virus particles (pink). **(C)** zoomed in 2D slices through 3D tomograms of virus particles 1–3 from (A) (10-slices averaged). **(D)** 2D slice through a 3D tomogram of a *N. marumarumayae* cell (10-slices). **(E)** Segmented volume D, showing features coloured as in (B) with the interacting virus particles shown in (orange). **(F)** zoomed in 2D slices through 3D tomograms of virus particles 1–5 from (D) (10-slices). Scale bars represent 100 nm.

In the first case, the siphovirus morphology is typical of many *Caudoviricetes*^33^. The capsid 51.1 ± 1.72 nm (n=5) in diameter appears to be icosahedral in shape, a tail 7.6 ± 0.9 nm (n=5) in width, extends 56.3 ± 6.2 nm (n=5) from the base of the capsid toward *N. marumarumayae* cell membrane (Fig. 2A and Extended Data Fig. 1A). Sheathed tails are typically 20–34 nm wide^34,35^, suggesting that this tail was not sheathed and is thus representative of siphovirus morphologies. The *N. marumarumayae* cell membrane density and irregular proteinaceous outer layer^20^ made visualising protein densities near the cell surface difficult (Fig. 2 and Supplementary Video 1). It is possible that this phage has either long and or short tail fibres that are not observable.

The second morphology is distinctively unique (Fig. 2B and Extended Data Fig. 1B, C). The capsid appeared prolate in shape, however, the top and bottom may also be conical structures rather than curved, as they appeared straight in some views – a structure resembling spindle virus capsids^36^ (Fig. 2B and Supplementary Video 1). The capsid had a width of 28.1 ± 2.1 (n=5) and height of 57.3 ± 11.5 nm (n=5), a unique component of this morphology was four distinct collar densities beneath the capsid, organised into two pairs, each pair was connected to the capsid by a weaker density (Fig. 2B and Extended Data Fig. 1C). Each collar protein pair contained a larger density proximal to the capsid, 4.4 nm across, and a smaller density distal to the capsid, 3.9 nm across. The length of the weaker density that connects the capsid to the collar proteins was 6.2 nm; a cross-section view shows that there are six pairs in total (Fig. 2B and Extended Data Fig. 1C). Finally, at least two fibres were seen extending 41.0 nm ± 3.9 nm (n=5) towards the cell surface, these fibres were too thin to accurately measure (Fig. 2B and Extended Data Fig. 1B). However, they are similar in length and thickness to the outer fibres of the host, suggestive of a potential co-ordination of the host fibre and interaction with the virus.

Spindle-shape morphologies have been previously predicted from metagenomic analyses of Asgard archaeal viruses, specifically Munnin virus^12^ and members of *Wyrdvirus* spp.^13^. Our observations therefore provide direct visual evidence linking this morphotype to an Asgard archaeal host. Notably, both virus morphotypes were observed not only on the surface of the main cell body, both also on the surface of the envelope vesicles protruding from the *N. marumarumayae* cells^20^ (Fig. 2 and Supplementary Video 1). No cells were observed simultaneously associated with both viral morphotypes. Due to the complexity of these cultures and the challenges of culture enrichment, lower abundance of *N. marumarumayae* cells on cryo-EM grids, and difficulty of identifying VLPs before tomogram reconstruction, we only observed VLP interaction with *N. marumarumayae* cells in two tomograms (multiple incidents in each tomogram), despite extensive screening.

### Correspondence of *N*. *marumarumayae* viral genomes to cryo-ET

To identify viral genomes corresponding to the viruses observed by cryo-ET, we analysed short- and long-read metagenomics datasets generated from a collection of enrichment cultures spanning several years (Supplementary Fig. 1)^20^. Three viruses were consistently associated with *N. marumarumayae* based on a combination of host-linked CRISPR spacers, Hi-C and genomic predictions. These viruses were designated *N. marumarumayae* viruses 1–3 (NMV1–NMV3) (Fig. 3A and Supplementary Table 1).

**Figure 3.**
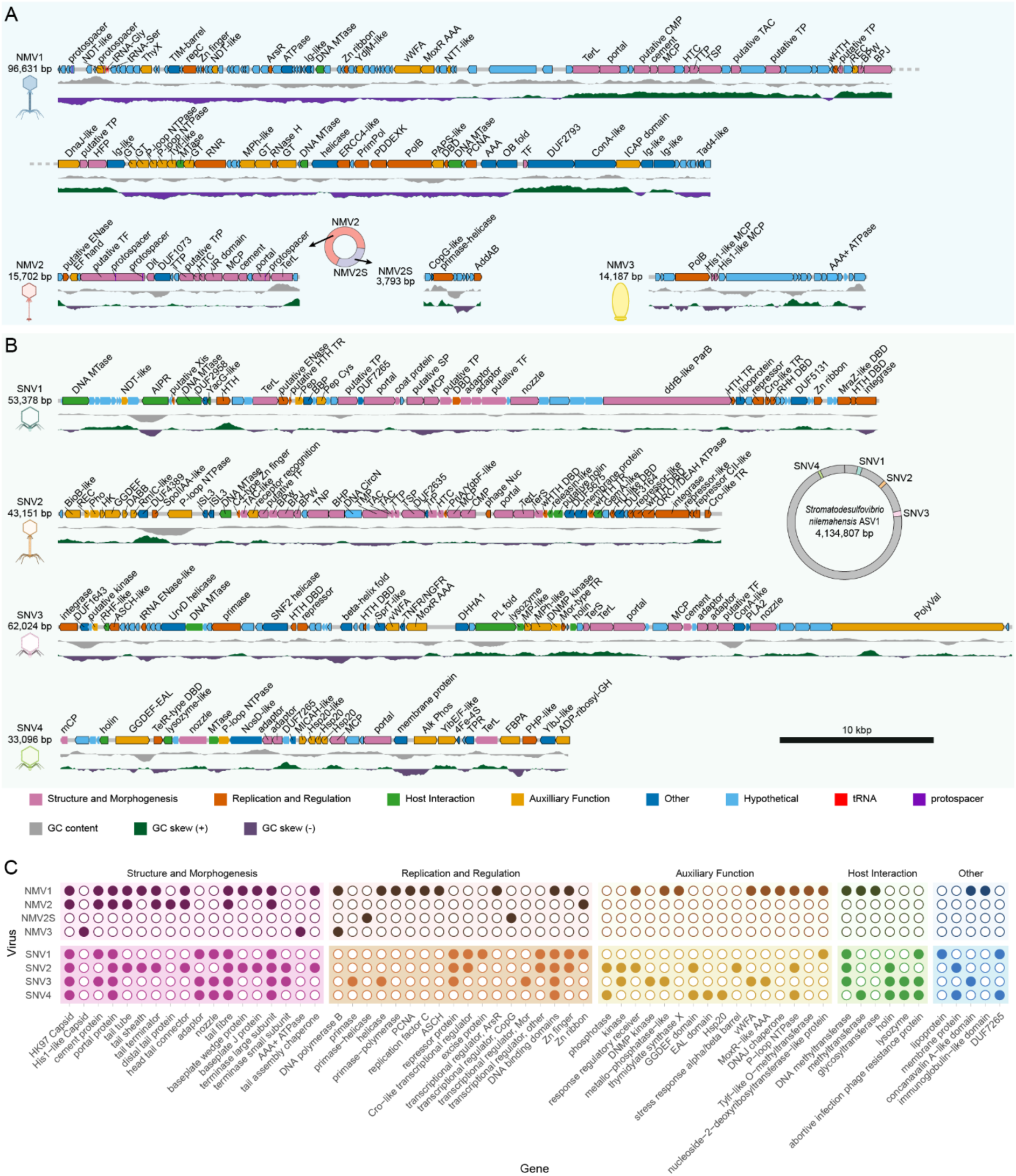
Overview of enrichment culture (pro)viruses and their functional annotations. Genome maps of NMVs **(A)** and SNVs **(B)** show ORFs (colour-coded by functional group), protospacers, and tRNAs. GC content and GC skew tracks are shown below the maps, with predicted morphotype icons to the left. Circular schematics show: A) the genomic integration of NMV2 and NMV2S, and B) the location of the SNVs within the *S. nilemahensis* Desulfo-ASV1’s genome. **(C)** Summary of functional annotations per provirus, presented as presence/absence (filled/empty circles, respectively).

NMV1 was assembled as a circular genome of 96.6 kbp with terminal repeats, indicating a closed complete genome. Two protospacers from an *N. marumarumayae* CRISPR system matched NMV1, targeting genes encoding a hypothetical protein, and a nucleoside-2-deoxyribosyltransferase-like protein (Fig. 3A and Supplementary Table 2). NMV2 was also targeted by the *N. marumarumayae* CRISPR system, with three unique protospacers matching a portal protein, and putative tail fibres (Fig. 3A and Supplementary Table 2). In contrast, NMV3 was recovered as a 14.1 kbp linear contig (Fig. 3A), which is similar to previously characterised dsDNA archaeal viruses^37^. While no CRISPR spacers were identified targeting NMV3, Hi-C analysis revealed strong virus-host contact linkages between NMV3 and *N. marumarumayae* (Supplementary Table 3).

Given cryo-ET revealed only two virus morphotypes associated with *N. marumarumayae*, we next aimed to link viral genomes to observed VLP structures using genomic annotations and structural modelling of predicted viral proteins (Extended Data Fig. 2 and Supplementary Figs. 2–5). First, both NMV1 and NMV2 had hallmark proteins characteristic of head-tail viruses from the class *Caudoviricetes*, including a HK97-fold^38^ (only identified in NMV2 via structural modelling), major capsid protein (MCP), portal proteins and terminase large subunits (Fig. 3, Extended Data Fig. 2, Supplementary Table 1 and Supplementary Figs. 2 and 3) By contrast, NMV3 encodes MCPs specific to archaeal spindle viruses^39,40^, specifically quite similar to haloarchaeal virus His1 (family *Halspiviridae*)^32^ (Fig. 3A and Extended Data Fig. 2). Structural modelling and electrostatics analysis of NMV3 major capsid protein further identified hydrophobicity^39^ and *N*-glycan regions^36^ characteristic of spindle virus capsids (Extended Data Fig. 2E). Additionally, NMV3 encodes an AAA+ ATPase, where remote homology searches revealed similarity to *Tubulavirales* filamentous phage assembly proteins (Fig. 3 and Supplementary Table 4). The same annotation occurs in the AAA+ ATPase of predicted Asgard spindle Wyrdviruses, where they may play a role in Asgard spindle virus egress. Together, these structures strongly suggest that NMV3 corresponds to the spindle-shaped virions observed by cryo-ET.

Alongside the HK97 MCP, portal proteins and a terminase large subunit, NMV1 encodes additional hallmark proteins of head-tail bacteriophages including baseplate J and wedge proteins^27,28,41,42^. Structural modelling of monomeric and homomeric multimers further identified a cement protein, head tail connector, tail terminator, capsid fibre protein, tail tube, tail sheath, distal tail protein, and other tail/baseplate components (Fig. 3A, Supplementary Table 4 and Supplementary Fig. 2). These genes predominantly clustered within a virus morphogenesis module spanning 33,476–57,406 bp (Fig. 3A), with the presence of a contractile tail, inferred from these structural features, a hallmark of myoviruses^33^ (Extended Data Fig. 2A,B). In contrast, NMV2 lacks a predicted tail sheath but encodes for a terminase large subunit, tail protein, portal protein and MCP^27,28,41,42^. Structural modelling identified a tail fibre protein (Fig. 3A and Supplementary Fig. 3). However, given the absence of a tail sheath, NMV2 is predicted to have a non-contractile tail consistent with a siphovirus morphology^33^. Additionally, high confidence trimeric models of the NMV2 tail tube closely matched the dimensions of the tail width of the siphovirus morphotype that was associated with the *N. marumarumayae* cells in the cryo-ET (Fig. 2, Extended Data Figs. 1A and 2A, B), suggesting that the head-tail virus associated with *N. marumarumayae* cells in the cryo-ET is likely to be NMV2 (Fig. 2, Extended Data Figs. 1A and 2A, B).

### A putative virus satellite is associated with NMV2

NMV2 was initially recovered from metagenomes generated in 2023 (Supplementary Fig. 1) as a clean circular 19.6 kbp genome with uniform read coverage (Fig. 4A). However, analysis of cultures sequenced from 2024 onwards (Supplementary Fig. 1) consistently resolved this element into two distinct components: a 15.7 kbp intact linear core contig corresponding to the core NMV2 virus and a 3.8 kbp circular element present at substantially higher coverage (Figs. 3 and 4a) indicating higher copy numbers. Interestingly, long reads that mapped to the circular component displayed a variety of structural variants including concatemers with some reaching up to15.2 kbp (4-mer) and longer, similar in length to the NMV2 core linear genome (Fig. 4C). While such concatemers could be a result of rolling circle replication^43^, concatemers have been reported in satellites as a mechanism to elongate their genome to fit helper virus capsid^44^ suggesting that this circular genome may be a satellite. Although most ORFs in the high-coverage circular element lacked functional annotation, several ORFs exhibited homology to proteins associated with plasmid-derived satellite elements^44,45^ of haloarchaea and nanohaloarchaea^46^, including an Arc-like regulatory protein, a helicase-primase, and a Cas4-like nuclease (Fig. 3A and Supplementary Table 4). Together this evidence supports similarity to a satellite and therefore, we designate this circular element as NMV2S and hypothesise that it functions a putative satellite that is dependent on NMV2 as a helper virus^47^.

**Figure 4.**
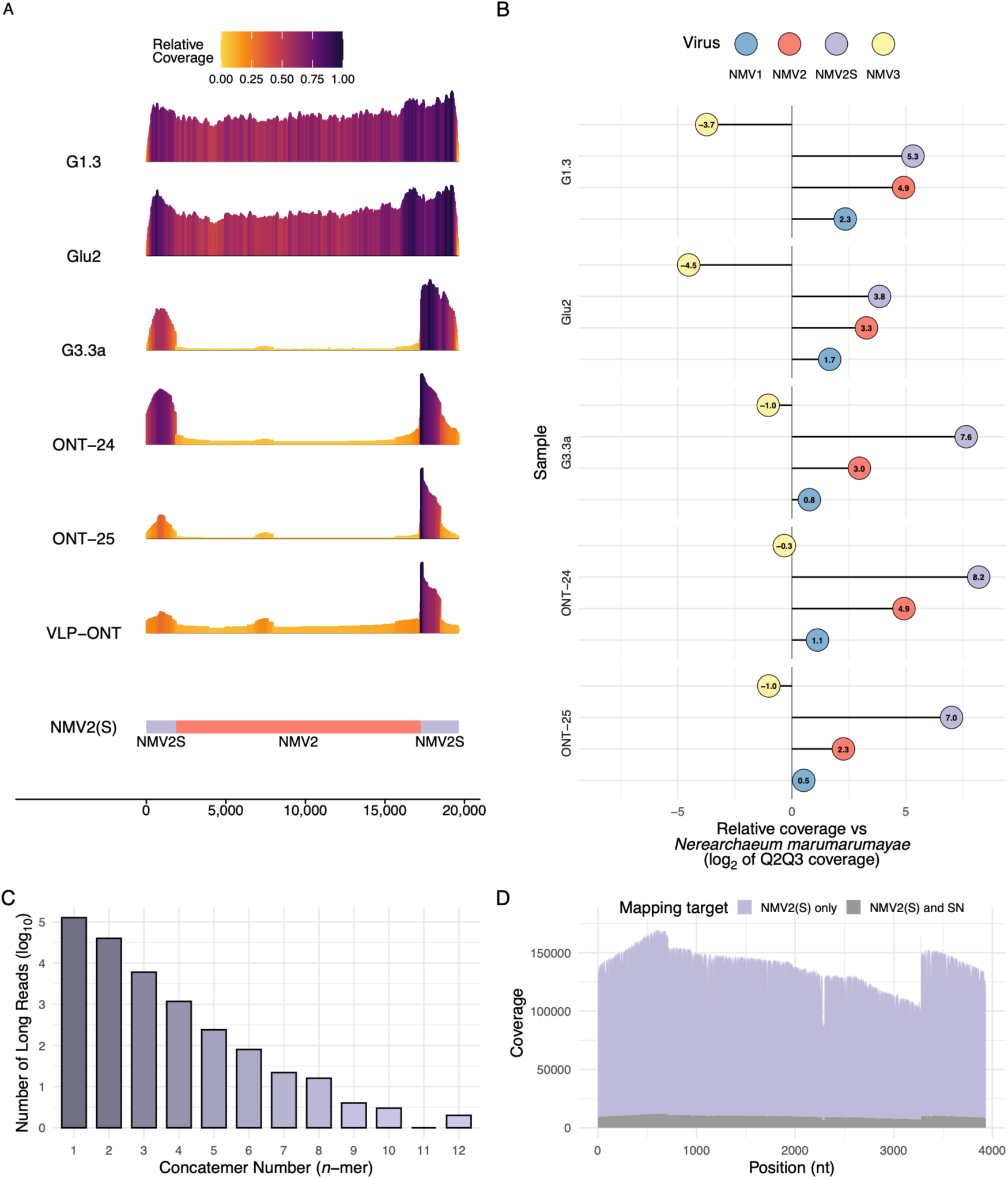
Unique replication dynamics of NMVs and the putative satellite element NMV2S. **(A)** Relative coverage profiles of NMV2 and NMV2S across metagenomic samples collected over time. **(B)** Log_2_ transformed relative coverage of NMVs with respect to *N. marumarumayae* Loki-ASV2 across metagenomes. **(C)** Length distribution of NMV2S concatemers recovered from individual long reads. **(D)** Coverage distribution of long reads mapping to NMV2S and to *Stromatodesulfovibrio nilemahensis* Desulfo-ASV1, highlighting reads shared between the satellite element and the bacterial genome.

To further investigate the potential replication mechanism of NMV2S, long reads from the viral fraction of the enrichment cultures (Supplementary Fig. 1) were generated and mapped to the NMV2 and NMV2S genomes. Assembly of the mapped reads recovered not only the viral sequences but also most of the closed *S. nilemahensis* genome. Mapping these reads back to the *S. nilemahensis* genome showed uniform coverage across the chromosome. Notably, a subset of reads that mapped to NMV2S also aligned to the *S. nilemahensis* genome, suggesting that some long sequencing reads span both viral and host sequences (Fig. 4D).

Viruses associated with *N. marumarumayae* exhibit distinct replication dynamics

Transcriptional activity was detected across all three NMV genomes (Fig. 3A and Supplementary Table 5), indicating that each virus was actively replicating within the enrichment cultures. Typical of viruses with larger genomes^37,48^, NMV1 encodes many core DNA replication genes, suggesting the capability of semi-autonomous genome replication, namely DNA polymerase B, replication factor C small subunit, bifunctional DNA primase/polymerase, helicase and ribonuclease H (Fig. 3A and Supplementary Table 4). Many of the listed NMV1 replication genes were predicted to be encoded by previously described Asgard archaeal viruses: Skoll, Fenrir, Nidhogg and Ratatoskr^14^.

NMV1 additionally encodes multiple DNA-binding proteins and endonucleases, including an ERCC4-like endonuclease, putative endonucleases with PD-(D/E)XK domains, as well as an ArsR type transcriptional regulator like domain, which may be involved in DNA replication and repair, or regulating transcription (Fig. 3A and Supplementary Table 4). Similar endonucleases, including an ERCC4-like endonuclease, have been reported in Asgard archaeal viruses, and were proposed to play a role in DNA repair^14^.

In contrast, the DNA replication mechanism of NMV2 is unclear, as the virus lacks genes indicative of viral replication, thus NMV2 likely relies on host replication machinery for viral replication (Fig. 3A and Supplementary Table 4). The associated satellite of NMV2, NMM2S, encodes a Cas4-like nuclease found in a number of bacterial and archaeal viruses^49,50^. Given the recombinase function of Cas4 enzyme nuclease domains^51^, NMV2S may utilise the Cas4-like nuclease for recombination and integrate within the NMV2 genome (Figs. 3a and 4a). NMV2S may assist in replication whilst integrated within NMV2 through the encoded helicase-primase, otherwise it is likely NMV2 largely relies on host replication machinery. Recruiting host replication machinery is typical of small archaeal virus genomes, including the Asgard archaeal virus genus ‘Verdandivirus’^13^. Like NMV2, NMV3 may have minimal capacity for autonomous replication given that it only encodes its own DNA polymerase (Fig. 3A and Supplementary Table 4).

Despite persistent presence of all three NMVs in enrichment cultures detected via PCR (Supplementary Fig. 6 and Supplementary Table 6), genes indicative of typical viral lifestyles, such as lysins or integrases, were not detected (Fig. 3A and Supplementary Table 4). Five Asgard archaeal enrichment metagenomes (Supplementary Fig. 1) were mapped back to the NMVs and their host (Fig. 4B) revealing that NMV abundances were consistently higher than their hosts, with exception to NMV3 with abundance consistently lower than the host (Fig. 4B). Additionally, no significant (p value < 0.05) correlations were observed between virus-host abundances (Supplementary Table 7). This data indicates that NMVs do not follow classical host-coupled replication dynamics and instead they may follow a more complex lifestyle^52^. In contrast, NMV1 and NMV2 exhibited significant (p value < 0.05) positive correlations across all samples, as well as NMV2S and NMV3 (Supplementary Table 7), potentially indicating shared optimal replication conditions or co-infection dynamics.

### *Stromatodesulfovibrio*-associated proviruses exhibit differing levels of activity and genome organisation

Four putative viruses associated with *S. nilemahensis* were identified as proviruses integrated within the bacterial host genome (SNV1–4; Fig. 3B and Supplementary Table 1). Integrated proviruses can be actively replicating, switching to a lytic lifestyle, or remain lysogenic even losing the ability to produce phage particles (inactive), known as legacy or cryptic phages^53^. Therefore, we assessed proviral replication activity by sequencing the viral fraction, then read-coverage analysis and prediction of attachment sites. Sharp shifts in coverage in the *S. nilemahensis* genome were observed at the predicted boundaries for SNV1 and SNV3 (Extended Data Fig. 3), indicating increased copy numbers relative to the host genome. Additionally, assembly of these mapped reads resulted in circularised genomes for SNV1 and SNV3, providing strong evidence that they are both actively replicating viruses. Although no attachment sites were predicted for both these genomes, SNV3 was integrated between two host tRNAs (Extended Data Fig. 3), typical of many prophages^54^.

Two sets of alternative attachment sites were identified in SNV2. The first set of predicted attachment sites may represent original viral boundaries, and subsequent recombination events may have expanded these boundaries to incorporate the auxiliary viral genes, with auxiliary viral gene acquisition through HGT well documented in diverse environments^55,56^. Both predicted *attR* sites were upstream of a host tRNA and in proximity, with a drop in read coverage observed upstream of the host tRNA and downstream of the predicted *attR* sites. However, no coverage changes were detected at either of the predicted *attL* sites (Extended Data Fig. 3). The first pair of attachment sites surrounded the ‘viral component’ of SNV2, containing only the core viral genes (Fig. 3B and Extended Data Fig. 3) and corresponding with a positive GC skew, suggesting a proviral region. The second pair of attachment sites matched the original predicted viral boundaries and coincided with a drop in observed read coverage. This attachment site pair encoded a module of auxiliary viral genes and displayed a drastic change in GC content, typical features of MGEs or HGT events^57^ (Fig. 3B and Extended Data Fig. 3).

The organization of this module bares similarity to known bacterial and signalling architectures that modulate biofilm formation^58,59^, consistent with the established ability of lysogenic phages to carry biofilm-associated auxiliary genes^60–62^. The auxiliary viral gene cluster began with a biotin synthase-like radical SAM enzyme, followed by a response regulator receiver domain, a phosphatase, and a kinase—typical of two component regulatory systems^63^. Adjacent to this module was a diguanylate cyclase and a stress response A/B barrel protein. Expression of all genes in this cluster was detected except for the response regulator receiver and phosphatase (Supplementary Tables 4 and 5). Horizontal acquisition of this cluster would account for the asymmetric coverage pattern observed: the left boundary shows only subtle fluctuations, whereas the right boundary displays a sharper transition where the two predicted *attR* sites lie in close proximity (Extended Data Fig. 3).

In contrast, there were no observable changes in coverage, and no clear indicators of viral boundaries for SNV4. Together with the lack of an integrase and its mosaic genome, our observations suggest SNV4 is a legacy virus (Fig. 3B and Extended Data Fig. 3). However, an expressed cluster of heat shock family (HSP20)^64^ and an expressed bifunctional diguanylate cyclase/phosphodiesterase proteins with functions in biofilm formation and stress response were identified between viral hallmark genes^65^ (Fig. 3B and Supplementary Tables 4 and 5). Together with the signalling cluster encoded in SNV2, a propensity for SNVs to encode or replicate in association with regions of these functions is proposed, which may modulate these functions in the host.

### *Stromatodesulfovibrio* proviruses exhibit diverse lifestyles and virion structures

In contrast to NMVs and consistent with their temperate lifestyles, transcriptional expression was detected for only some SNV ORFs^66^ (Fig. 3B and Supplementary Table 5). Unlike the Asgard archaeal viruses, the bacterial SNVs encoded transcriptionally expressed hallmark genes associated with lysogenic lifestyles, including recombinases and integrases (Fig. 3B,C and Supplementary Table 5). SNV2–4 encoded genes consistent with cell lysis, including putative holin proteins and lysozymes, with expression only observed for SNV4. In contrast, SNV1–3 encoded transcriptionally expressed repressor proteins, likely involved in maintaining a lysogenic state^67^ (Fig. 3B,C and Supplementary Tables 4 and 5). SNVs also encoded for proteins with antagonistic functions that potentially modify or hijack host replication, e.g., homologues of proteins with gyrase inhibitor functions, or YacG-like and GemA-like, respectively in SNV1 and SNV2 (Fig. 3B). Members of both of these families inhibit host DNA gyrase activity and may promote relaxation of the host’s genome, thus assisting phage replication^68,69^.

All four SNVs encode hallmark proteins of the order *Caudoviricetes*, including terminase subunits, HK97-fold MCP, and a portal protein (Fig. 3B,C and Supplementary Table 4)^27,28,41,42^ . Despite these shared features, structural modelling predicted differing viral morphologies (Fig. 3B and Supplementary Figs. 7–10). SNV2 is predicted to form a contractile tail morphology similar to NMV1, with structural modelling revealing a tail sheath, tail tube, tape measure protein, tail terminator protein, and other tail/baseplate components, (Fig. 3 and Supplementary Fig. 8) within a morphogenesis module (11,822–31,509 bp), including both large and small subunits of the DNA-packaging terminase (Fig. 3B). In contrast, SNV1, SNV3, and SNV4 encode tail adaptor and nozzle proteins indicative of a tail-less morphology typical of podoviruses^33^ (Fig. 3B,C and Supplementary Figs. 7–10). SNV1 and SNV3 encode their structural proteins within clustered morphogenesis modules that include packaging terminases (Fig. 3B, Supplementary Table 4). SNV4 structural genes were dispersed throughout the proviral genome, which may reflect shorter genome length or genetic transfer events that have disrupted gene clustering (Fig. 3B and Supplementary Fig. 10). Notably, SNV4 encodes only the large terminase subunit, with no detectable small subunit (Fig. 3B).

Typical of prophages, SNVs predominantly utilise host replication machinery for viral replication^70^ (Fig. 3B). SNV3 was the only provirus to encode replication machinery, including proteins with helicase and primase functions, all of which were transcriptionally expressed (Fig. 3B,C and Supplementary Tables 4 and 5). Additionally, an expressed gene encoded a protein belonging to the tRNA endonuclease-like domain superfamily (IPR011856), which may be involved in replication, such as resolvase and restriction endonuclease activity^71^ (Fig. 3B). SNV3 also encodes a transcriptionally expressed homologue of ASCH domain, which may play a role in RNA binding and regulation of expression^72^, or detection of modified bases as part of defence systems^73^ (Fig. 3B, Supplementary Table 5). Across all SNVs, numerous uncharacterised proteins with predicted DNA binding, HTH or Zn finger domains were identified, which may play a role in transcriptional regulation or recruiting host replication and transcriptional machinery^13^ (Fig. 3B, Supplementary Table 4). Most of these proteins in SNV1 and SNV3 were transcriptionally expressed, whilst SNV2 only had a Hu-like DNA binding domain and a winged HTH DNA binding domain expressed (Supplementary Table 5). Lastly, only one gene of uncharacterised DNA binding domain protein was expressed in SNV4 (Fig. 3B, Supplementary Table 5). While beyond the scope of this study, follow up studies can examine viral gene expression under different conditions to clarify the role of all ORFs identified.

### Classification reveals all viruses as novel orders

Taxonomic classification based on protein network clustering performed by vConTACT3, (Fig. 5) assigned all NMVs and SNVs identified as novel orders, genera and species (Supplementary Tables 1 and 8). NMV1 is designated as belonging to a novel order within *Caudoviricetes*. This order also contains scaffolds representing previously described Asgard viruses: Fenrir viruses^14^ (Supplementary Tables 8 and 9). However, Fenrir viruses and NMV1 each belong to distinct novel families. We designate NMV1 as belonging to a new genus, *Brokkivirus*, and as the only current species within the genus; *Brokkivirus salinus* (Fig. 5, Supplementary Tables 1 and 8–10). NMV2 was unrelated to all other viruses except for two viral scaffolds from metagenomes of Antarctic hypersaline lake Ace Lake^74^ (Fig. 5, Supplementary Tables 8 and 10). These viruses are related to NMV2 at the family level but from different genera, with Ace Lake viruses sharing the same genus (Fig. 5, Supplementary Table 8). Therefore, we designate NMV2 to a novel genus and species; *Juurluvirus versutum* (Supplementary Table 1). Although not assigned a realm at the sequence level by protein clustering, NMV2 still is characterised as *Caudoviricetes* given multiple hallmarks including the HK97-fold MCP (Fig. 3, Extended Data Fig. 2D). NMV3 was also not assigned a realm by vConTACT3 and represents a novel order shared with viral scaffolds attained from JGI blast search of NMV3 capsids (Fig. 5, Supplementary Tables 8–10, Methods). The order diverges into two novel families, with NMV3 being a novel species in its own genus in one of these families, which we designate as *Logivirus ellipsoideus* (Fig. 5, Supplementary Table 1).

**Figure 5.**
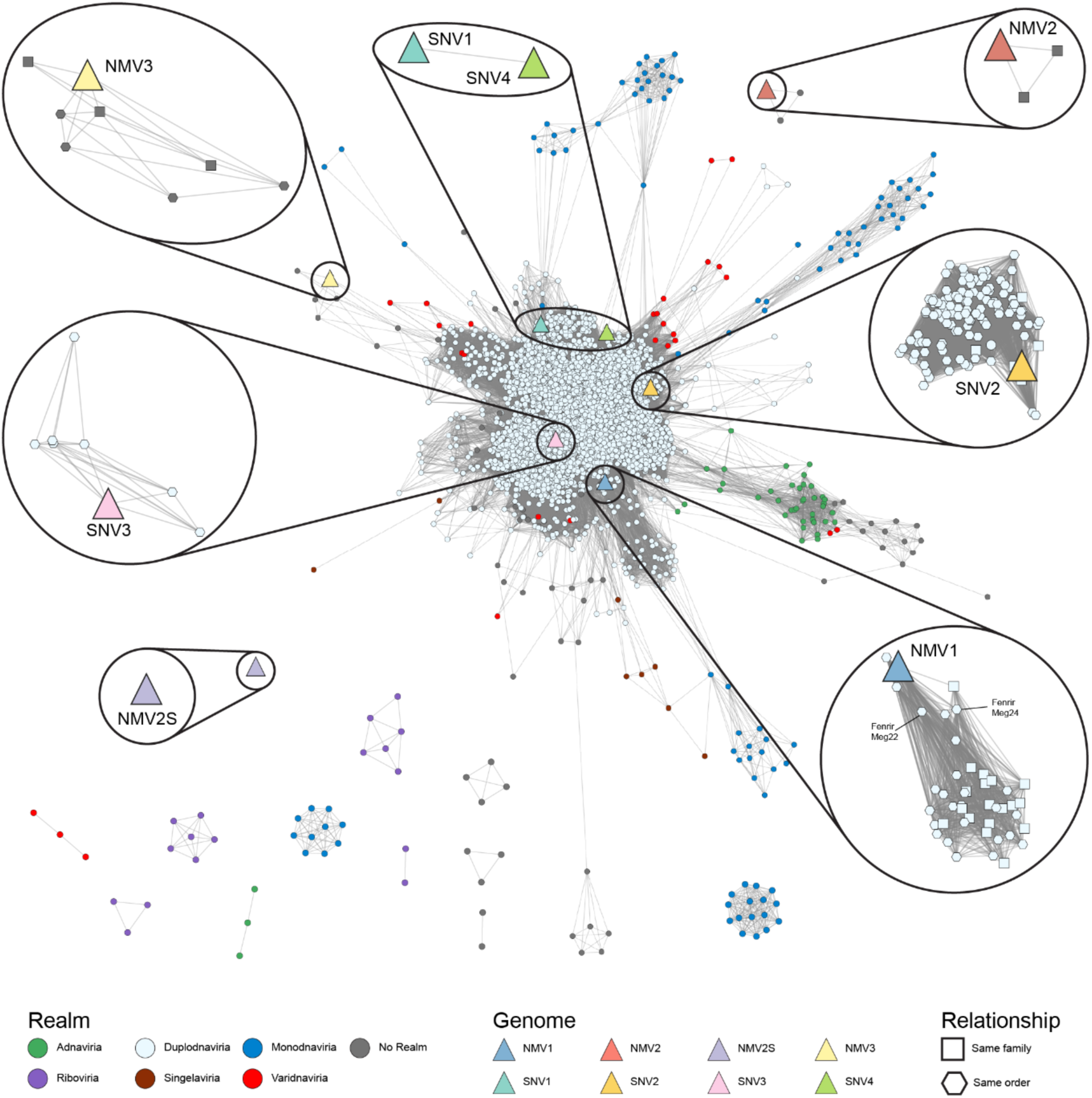
Protein-sharing cluster network analysis of NMVs and SNVs generated using vConTACT3. Reference virus members are coloured by realm, whereas NMVs and SNVs are represented as triangles. Insets show enlarged views of the order-level subnetworks containing viruses identified in this study. Node shapes indicate the taxonomic relationship of reference viruses to NMVs and SNVs.

SNV1 and SNV4 form a novel order, as the only members of this order, within *Caudoviricetes* but are predicted as belonging to separate families (Fig. 5 and Supplementary Table 8). Given that SNV4 is likely a legacy virus, it is possibly a remnant of an SNV1 infection. We have assigned SNV1 the name *Stromatovirus australiense* (Supplementary Table 1). SNV2 forms a novel order with already described families including the family *Casjensviridae*^75^ (Fig. 5 and Supplementary Table 8). Other families in this order contain already described genera which have not yet been assigned a family. SNV2 is in a novel family within this order which would include the *Muvirus* genus^74^ (Fig. 5 and Supplementary Table 8). SNV2 is the only species within a novel genus in this family, which we designate as *Firmivirus cohaerens* (Supplementary Table 1). SNV3 forms a novel order which includes a novel family of previously described *Clostridium perfringens* phages^76,77^, with no ICTV assigned order. SNV3 is the only member of its own novel family, therefore also forming a novel genus and species (Fig. 5 and Supplementary Table 8). We have assigned the name *Stronivirus nilemahense* to the species of SNV3 (Supplementary Table 1). NMV2S was classified as a singleton with no realm annotated (Fig. 5). We have named NMV2S *Trinitisatellite surturii* (Supplementary Table 8). Naming and designations of viruses described in the present study are summarised in Supplementary Table 1.

## Discussion

Interactions between ancestral Asgard archaea and bacterial partners are central to leading models of eukaryogenesis, yet the contribution of viruses to this pivotal evolutionary process has remained largely unexplored. Here we provide the first integrated characterisation of viruses associated with an Asgard archaeal enrichment culture and the first direct visualisation of viruses attached to Asgard archaeal cells. By combining cryo-ET, long-read sequencing and Hi-C, we identified three viruses associated with *N. marumarumayae*, including a spindle-shaped morphotype, four proviruses associated with *S. nilemahensis*, and a putative satellite element linked to an Asgard archaeal virus. Together, these findings reveal an unexpected level of complexity within an archaeal–bacterial co-culture relevant to understanding interactions among lineages implicated in models of early eukaryotic evolution.

The three *N. marumarumayae* viruses (NMV1–3) differed substantially in morphology, gene content, and predicted replication strategies, indicating that multiple viral lifestyles may co-exist in a stable Asgard archaeal host population (Fig. 3). Despite persistent detection across enrichment cultures (Supplementary Fig. 4), fluctuations in viral abundance were not coupled to host abundance (Fig. 4A). This pattern deviates from classical virus–host systems^52^, which is more consistent with chronic, low-impact or otherwise non-lytic infection strategies that permit sustained viral production without measurable host decline^52,78,79^. Whilst described spindle viruses all establish chronic infection, all members of *Caudoviricetes* are known to establish a lytic state at the end of their infection cycle^13^, however the Asgard archaeal viruses observed may only infect a small proportion of the host population^80^ or regulate infection stages and stall lysis^81^. Although viral particles were observed on the surface of host cells, no definitive entry events or co-infections were captured (Fig. 2), potentially suggesting tightly regulated or prolonged early infection stages. Together these results suggest the observed diverse viral lifestyles may enable a persistent, low impact infection of their Asgard archaeal host, raising important questions about how multiple viruses partition resources, avoid direct competition, or even interact cooperatively within a single slow growing host.

One of our most intriguing observations was the association of viruses with membrane vesicles protruding from the *N. marumarumayae* cell body^20^ (Fig. 2). The biological role of these structures remains unresolved. They may function as decoys, as part of a host immune response, that divert viral attachment away from the main cell body as in some bacterial species^82^. Alternatively, they may be a direct result of viral propagation – with Archaeal viruses previously shown to manipulate host cells and exit through a budding-like mechanism^31,36,78^. Regardless of their function, the increase in surface area resulting from these vesicles may “trap” viral particles, with viral particle entrapment in biofilm ecosystems important in biofilm formation and stability^83–85^. Although further experimentation is required, these observations establish a previously unrecognised connection between Asgard archaeal viruses and extracellular membrane structures.

The discovery of NMV2S, a putative satellite associated with NMV2, further expands the known complexities of Asgard –associated viral systems. Virus satellites are important regulators of viral ecology because they alter helper-virus propagation^86^, replication dynamics and transmission^87^. Given NMV2S abundance was similar or substantially higher than NMV2 (Fig. 4), NMV2S likely outcompetes NMV2^46^ and may be contributing to a stable host population, despite persistent viral infection. The identification of this viral satellite will be key in future studies of helper–satellite dynamics in Asgard archaeal systems.

Strikingly, long read assemblies revealed genomic interactions between NMV2S and *S. nilemahensis* (Fig. 4). Although the mechanism underlying these interactions remains unresolved, they provide evidence of an Asgard archaeal associated mobile genetic element connecting with a bacterial partner in co-culture. Cross-domain viral interactions are proposed as a viral adaptation strategy which may mediate inter-domain gene flow, with such interactions previously documented in microbial mat communities^26^. Additionally, previous observations of a nanotube from *S. nilemahensis* to *N. marumarumayae* with unknown function^20^ suggest a potential mechanism for the exchange of small genomic elements such as NMV2S^88^. While experimental confirmation of host range expansion and transfer mechanisms will be required, our findings provide a basis for investigating cross-domain viral interactions in similar enrichment culture systems.

The *S. nilemahensis* proviruses provide additional evidence that viruses may influence community-level processes. These bacterial proviruses encoded auxiliary functions associated with stress response and biofilm formation (Fig. 3). Whether these genes affect biofilm formation in *S. nilemahensis,* as suggested in previously described systems^60–62^, remains to be experimentally determined, but their repeated detection suggests they could be an important feature not only in the enrichment cultures, but also microbial mat ecosystems, given biofilms are defining features of microbial mats.

Collectively, our findings support a framework for investigating how viruses influence archaeal–bacterial interactions with potential for future studies into the emergence of eukaryotes. We therefore propose a conceptual model (Fig. 6) in which diverse archaeal viral lifestyles enable persistent, low impact, infection of an Asgard archaeal host. Increased surface area of the Asgard archaeal host may be the result of a production of “decoys” as an immune response or a result of virus replication, a satellite–helper axis adds competitive and regulatory complexity, and bacterial-associated proviruses contribute auxiliary traits such as biofilm formation that may enhance physical interfaces among community members. More broadly these processes may increase opportunities for connections and interactions among archaeal, bacterial and viral partners within microbial mats^89^.

**Figure 6.**
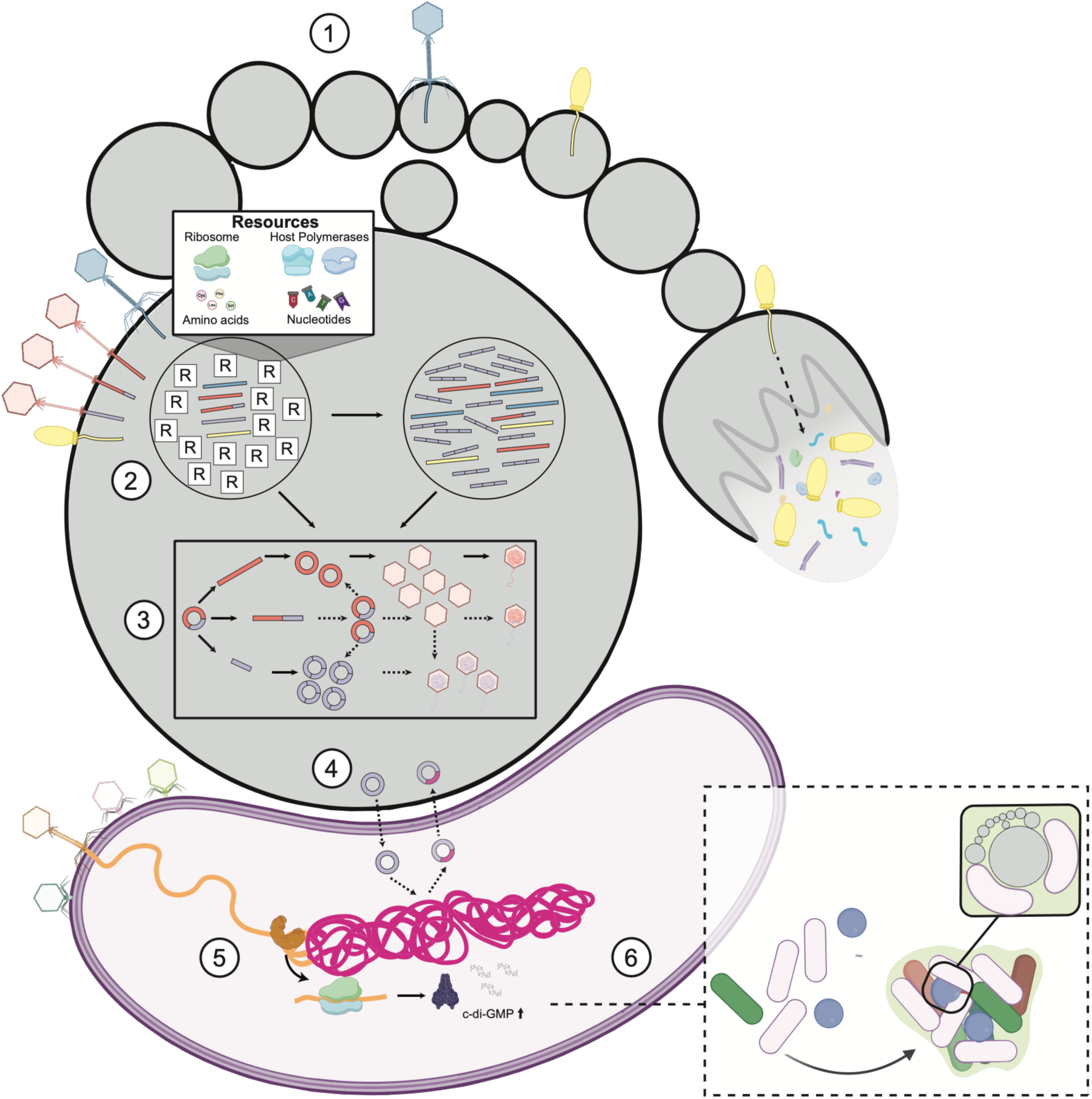
Conceptual model of proposed virus-host interactions inferred from the present study. Depicted are potential interactions between *Nerearchaeum marumarumayae, Stromatodesulfovibrio nilemahensis* and their associated viral elements. **1)** *N. marumarumayae* vesicles, increasing host surface area, may function (even inadvertently) as “decoys” as part of host immune response or generated directly from viral replication. **2)** NMV2S enters cell through NMV2 encoded virion particle and competes with other NMVs for resources for replication regulating the infection rate of NMVs. **3)** NMV2S outcompetes replication of NMV2 with higher genome copy numbers. NMV2S may package into NMV2 particles while remaining integrated in NMV2 genome or as concatemers. **4)** NMV2S genomic material enters *S. nilemahensis* through an HGT mechanism and potential genetic exchange between NMV2S and *S. nilemahensis* occurs. **5)** SNV auxiliary viral genes, including those with diguanylate cyclase activity, are integrated into host genome. Expression of these genes leads to increased cyclic-di-GMP in cell. **6)** increased cyclic-di-GMP in cell leads to lifestyle changes, including potential increased biofilm formation.

Although direct links to eukaryogenesis remain speculative, the systems described here provide a new framework for investigating how viruses influence Asgard archaeal–bacterial interactions which may have been key at different stages of the emergence of eukaryotes.

More broadly, our findings expand the known diversity of the Asgard archaeal virome and establish a foundation for exploring the role of viruses in microbial partnerships with potential relevance to early life/ eukaryogenesis.

## Methods

### Viral-like particles separation and imaging

For separating VLPs for imaging, enrichment cultures were centrifuged at 17,000 rpm for 30 min at 4 °C in Hitachi VX22N High Speed Centrifuge using R18A rotor. Supernatant was collected and stored at 4 °C before ultracentrifuging in Beckman Optima XPN-100 Ultracentrifuge using SW32 Ti rotor at 30,000 rpm at 4 °C for 4 h. Pellets were resuspended in high salts buffer (500 mM NaCl, 50 mM Tris-HCl, 5 mM MgCl_2_ at pH 8). Resuspended VLPs were probe-sonicated on ice using Qsonica Q500 Sonicator with a 1.6 mm microtip at 20% amplitude, for 3 × 10 s (2 s on 2 s off) with 10 s break between pulses. NEB Salt Active Nuclease was then added to a final concentration of 22 U/mL before being incubated at 1 h room temperature on a rotary shaker. Probe sonication was repeated, then samples were centrifuged in benchtop microcentrifuge (Eppendorf Centrifuge 5425) at max speed for 5 min. Supernatant was collected as the “soluble fraction” and spun at 50,000 rpm for 2 h at 4 °C in Beckman Optima XPN-100 Ultracentrifuge with SW55 Ti rotor. Pellets were resuspended in high salts buffer and prepared for transmission electron microscopy imaging (described below).

Carbon-coated 300 mesh copper grids were glow discharged in air using a PELCO easiGlow system (20 mA, 30 s, negative polarity) to create a hydrophilic surface. The carbon-side of each grid, held with self-closing forceps, was touched to a 5 µL drop of sample, and allowed to adsorb for 2 min. Excess sample was removed by gentle blotting with Whatman Grade 1 filter paper. Grids were then washed by sequentially touching the sample-side to four 5 µL drops of Milli-Q water, with blotting after each step. Grids were then negatively stained by touching the sample-side to a 4 µL drop of 2% (w/v) aqueous uranyl acetate and incubating for 1 min. Excess stain was removed by gentle blotting, and grids were air-dried in the dark.

Negative-stained grids were imaged using a Talos L120C transmission electron microscope (Thermo Fisher Scientific) operated at 120 kV with a LaB_6_ thermionic electron source. Brightfield TEM images were collected in microprobe mode, using EPU v3.11.0.9330 (Thermo Fisher Scientific). Images were recorded on a bottom-mounted Ceta-S camera at 45,000× nominal magnification as single frames with 1 x 1 binning (3.144 Å/pixel), 1 s exposure, and an electron dose of ∼13 e^−^ Å^−2^ per image. A nominal defocus range of -0.5 µm to -2.0 µm was applied during acquisition. A total of 15418 images were collected from 23 grid squares, selected for imaging based on appearance at low magnification.

All data processing was performed within the Scipion framework v3.8.3^90^. Images were imported using the pwem^90^ protocol with no phase flipping, spherical aberration set to 2.7 mm, amplitude contrast set to 0.1, and a sampling rate read directly from the image metadata. Contrast transfer function (CTF) parameters were estimated using CTFFIND^91–93^ implemented in cisTEM^94^. Particles were picked using the automatic particle picking workflow implemented in Xmipp3^95^. Briefly, 3122 manually picked particles were used to train an initial classifier. Output from this classifier was reviewed and corrected in a supervised learning phase, in which missed particles were added and false positives were removed. The retrained classifier was then applied to all remaining images, yielding 17,234 extracted particles. Due to the heterogeneity of virus-like particles and other particle types, extracted particles were visually assessed and a representative subset was selected to highlight the observed morphological diversity.

### Cryo-Electron Tomography

Samples from cultures designated as G2.24 and G3.3a were subjected to cryo-ET for high resolution analyses of viruses present. For freezing, samples were first mixed with 10 nm colloidal gold beads (Sigma-Aldrich, Australia), which were precoated with 1% BSA. The samples were then pipetted onto glow-discharged R2/2 Quantifoil holey carbon grids (Quantifoil Micro Tools GmbH, Jena, Germany), and excess liquid was removed by back blotting before plunging into liquid ethane. The entire freezing process was performed inside the Vitrobot chamber (FEI Thermo Fisher Scientific) under 100% humidity.

Grids were imaged using a Titan Krios G4 cryo-EM, operating at 300 kV acceleration voltage, and equipped with a Gatan energy filter and a K3 Summit direct detector. Tilt series images were collected in movie mode using a dose-symmetric scheme at pixel size, 3.39 Å, using Tomography 5 software v5.14 (Thermo Fisher Scientific) at a tilt range of −51° to 51° in 3° increments, with defocus values of -6 μm. Data were collection with a cumulative dose of ∼120 e^−^/Å² per tilt series. Frames were initially motion-corrected using MotionCor3^96^ and then aligned with the IMOD 5.1 software package^97^, integrated within ScipionTomo v3.0^98^. The aligned tilt series were then reconstructed at bin 4 using Tomo3D^99^. To enhance interpretability, tomograms were denoised using Cryo-CARE v3.2.0^100^.

To visualize and quantify tomograms, 3D volumes were segmented using Dragonfly software, v2022.2 (https://www.theobjects.com/dragonfly/index.html). Built-in filters were applied to improve clarity. This was followed by the manual segmentation of 20 slices from individual tomograms, which were then used as input for neural network training using the U-Net architecture with a 2.5D input dimension (7 slices)^101^. The trained model was subsequently used to segment the entire tomogram, with manual corrections implemented where necessary. Built-in functions were used to generate 2D images and 3D movies.

### DNA extraction

DNA was extracted from enrichment cultures using the NEB Monarch Genomic DNA Purification Kit following manufacturer’s instructions for purification from Gram-positive Bacteria and Archaea with the modifications previously described^20^: cells were collected by centrifugation of culture aliquots at 20000 × *g* for 10 min; after lysis and RNase treatment, sediment and cellular debris were separated from the lysate by centrifugation at 20000 × *g* for 10 min; and, supernatant was transferred to a new tube to be mixed with the gDNA binding buffer.

For extraction DNA of VLPs within “viral fraction”, enrichment cultures were centrifuged at 15,000 rcf for 30 min Supernatant was collected and stored at 4 °C. The pellets were collected and resuspended in a lysis salts buffer (300 mM NaCl, 50 mM Tris-HCl, 5 mM MgCl_2_ at pH 8, 0.1% Triton X, 50 U/ml benzonase). Sample was sonicated using Branson SFX250 Sonifier on ice with a 3mm tapered microtip at 45% amplitude, 6 × 10 s rounds with 10 s between each round of sonication. Sample was then mixed for 30 min at room temperature in rotary mix, before being centrifuged at 15,000 rcf for 30 min. Supernatant was collected and stored at 4 °C. Pellet was resuspended in a new lysis salts buffer (300 mM NaCl, 50 mM Tris-HCl, 5 mM MgCl_2_ at pH 8, 1% Triton X, benzonase) and sonicated again as above, mixed and centrifuged to collect lysate. Culture supernatant and lysate was collected and loaded onto a 25% sucrose cushion before ultracentrifugation overnight in Beckman Optima XE-90 Ultracentrifuge using SW28 Ti rotor at 25,000 rpm at 4 °C. Pellets were collected and resuspended in high salts buffer (300 mM NaCl, 50 mM Tris-HCl, 5 mM MgCl_2_ at pH 8) before performing Protein Qubit. Resuspended VLPs were then reconcentrated through ultracentrifugation in SW55 rotor at 50,000 rpm at 4 °C for 2 h and were resuspended in cold Tris-HCl 10mM pH 8 before immediately performing NEB gDNA extraction as above.

### DNA sequencing

Illumina shotgun metagenomic sequencing of sample SB19.G3.3a was performed on a NextSeq 1000 platform (Illumina) with a P1 flowcell using an Illumina DNA Prep library with 2×300 bp paired-end reads at the Ramaciotti Centre for Genomics. The reads were trimmed using the BBDuk module in BBMap v38.63 (https://bbmap.org) (ref=adapters,phix,lambda trimpolyg=1 qtrim=rl trimq=6). and subsequently assembled using metaSPAdes^102^ v3.15.5 with default settings.

For long read sequencing, three sets of pooled DNA extracts were prepared (≥200 ng DNA input per library): pool 1, containing DNA from VLP concentrates; pool 2, with DNA from the G5 lineage (114, 120 and 121); and, pool 3, with samples from the G4 lineage (62, 63, 64, 76). Libraries were prepared individually with the SQK-LSK114 kit as previously described^20^. Oxford Nanopore Technologies sequencing was performed in an ONT GridION platform for approximately 72 h per flowcell (R10.4.1) and two flowcells per pooled sample. The resulting data was basecalled offline with Dorado v0.9.1 (https://github.com/nanoporetech/dorado) and the corresponding super-accuracy model v5.0.0 in duplex mode (dna_r10.4.1_e8.2_400bps_sup@v5.0.0).

### Hi-C sequencing

Two ml aliquots of cultures were spun at 20,000 × *g*, with pellets washed in PBS and resuspended in PGShield™ prior to submission to Phase Genomics (Seattle, US) for library preparation and sequencing. The Hi-C library was created with the Phase Genomics ProxiMeta Hi-C kit v4.0 according to manufacturer’s protocol^103^. Separately, DNA was extracted from an aliquot of the same samples used for the Hi-C library preparation using the ZymoBIOMICS DNA Miniprep kit to generate a shotgun metagenomics library with the ProxiMeta library reagents. Both Hi-C and shotgun libraries were sequenced in an Illumina NovaSeq instrument as 2×150 bp paired-end reads.

### Metagenomic Assembly and Binning

Assembly of long-read data was performed with metaFlye v2.9.4-b1799^104^, reads provided with the --nano-corr option and --extra-params minimizer_window=10,repeat_graph_ovlp_divergence=0.005. Alternate assemblies of ONT-24 were performed with myloasm v0.1.0^105^ and hifiasm (ONT) v0.25.0-r726^106^, both with default parameters. Assembly graphs were manually evaluated to extract cMAGs and “promising subgraphs”. In the case of putative circular contigs observed in hifiasm or myloasm assemblies, these were compared to the Flye assembly graph in Bandage v0.8.1^107^ as they tend prematurely circularise contigs, esp. at lower coverages while Flye tends to keep them connected in the graph assembly. Reads were mapped against the selected subgraphs with minimap2 v2.29-r1283^108^ and reassembled as above, occasionally providing additional cMAGs. A final assembly with all the reads not mapping with a preliminary cMAG collection was performed and binned with TaxVAMB v4.1.4.dev150+g8fa3280^109^ and SemiBin2 v2.2.0^110^ (multi_easy_bin).

Short-read only assemblies (including the shotgun data from the Hi-C libraries) were performed on the reads not mapping the cMAG collection derived from the ONT assemblies. Bowtie2 v2.5.2^111^ was used for mapping and the unmapped reads extracted with SAMtools v1.13^112^ fastq command. Unmapped reads were assembled with MEGAHIT v1.2.9^113^ with the meta-sensitive preset. As per the ONT assemblies, TaxVAMB and SemiBin2 were used for binning.

Hi-C bins were generated by Phase Genomics bioinformatics pipeline (some software versions not available). Briefly, the shotgun reads were quality-trimmed and normalised with fastp^114^ and assembled with MEGAHIT and default parameters. Hi-C reads were aligned to the shotgun assembly using BWA-MEM v0.7.17-r1198-dirty^115^ with the -5SP option. PCR duplicated were flagged with SAMBLASTER v0.1.24^116^ for later removal. Non-primary and secondary alignments were removed with SAMtools view v1.13 (option -F 2304). Metagenome deconvolution was performed with ProxiMeta^117,118^. Resulting clusters were assessed for quality with CheckM^119^.

Long read bins, short read bins and Hi-C bins were all compared and dereplicated. In the case of long read bins overlapping with the either short read and/or Hi-C bins, they were consolidated by using quickmerge v0.3^120^, often improving the bin contiguity. All the redundant bins (including hybrid/merged) were also evaluated with CheckM2 v1.0.1^121^.

### Detecting Asgard archaeal viruses and host-matching

Putative viral contigs were determined using geNomad v1.11.0 ^122^ using default settings except score-calibration (--composition metagenome, --relaxed). Predicted viruses with a false discovery rate of 0.05 or lower were included in the viral dataset. CRISPR arrays in the complete *N. marumarumayae* MAG were predicted using the CRISPR-Detect^123^ webserver using previously described parameters^14^. *N. marumarumayae* CRISPR spacers were extracted and NCBI BLAST+ v2.16.0 ^124^ was employed to detect matches between spacers and the viral dataset (-task blastn-short -evalue 1e-5). A host match was accepted if there was a total of 2 bp or less that did not match across the entire spacer query.

To verify host matches, Hi-C reads were mapped to a curated collection of viruses and MAGs from enrichment cultures using BWA-MEM v0.7.17^115^ with the -5SP option. Unmapped reads, supplementary alignments, and secondary alignments were removed with SAMtools view v1.20^112^ (option -F 0×904). A normalized contact map of Hi-C linkages was produced using the NormCC normalization model of MetaCC v1.2.0^125^. Host-virus relationships were determined and filtered using a previously described method which utilizes Z-scores of normalized linkages^126^.

Trimmed reads from five different Asgard archaeal enrichment culture metagenomes were mapped to the *N. marumarumayae* viral genomes NMV1–3, NMV2S, and the *N. marumarumayae* genome using BBMap (https://bbmap.org). Subsequently, the mean Q2Q3 coverage for each genome and sample was formatted as a matrix and imported into RStudio (R v4.4.2). Spearman’s rank correlation coefficients and p-values were calculated for each pair and corrected for multiple test^127^ with the corr.test() function from the psych package v2.3.9 (https://CRAN.R-project.org/package=psych).

Long reads from sequenced virus fraction of enrichment cultures (see above) were mapped using minimap2^108^ v2.30 (-x lr:hq -c -p 0.5 -N 100) to NMV2S genome with alignment information written as a pairwise formatting (paf) file. A custom python script (github/link/to/script) was used to analyse the output PAF file to count the number of concatemers for each multiplicity (each “*n*-mer”) using the pandas library, run with Python v3.9.21 and pandas v2.2.3^128^. Prior to downstream analysis, alignments were strictly filtered to retain only high confidence matches, requiring a minimum mapping quality score (MAPQ) of ≥30. To minimize the inclusion of spurious mapping artifacts or negligible sequence fragments, individual alignment blocks were required to span a minimum alignment length (MIN_ALEN) of 1,000 bp. Because structural variants (such as large insertions or deletions) skew total read lengths, a step-wise, coordinate-based tracking algorithm was implemented to resolve the true multiplicity of individual long reads. For each unique query read ID (qname), all valid alignment blocks were grouped and sorted chronologically based on their physical start coordinates on the read (qstart). The physical length of the read was traversed sequentially. To prevent the double-counting of fragmented alignments or overlapping secondary mappings arising from mapping ambiguities, a secondary alignment block was only recognized as a distinct concatemer unit if its start coordinate did not overlap the preceding block’s end coordinate (qend) by more than 500 bp (MAX_OVERLAP). The final concatemer number (*n*) assigned to each individual read represented the total number of non-overlapping, high-confidence target intervals of >1,000 bp traversed along the reference. The resulting discrete *n*-mer frequencies were aggregated and exported to a structured tabular format for downstream distribution analysis.

Reads from viral fraction of enrichment cultures were mapped to NMVS2 genome using minimap2 v2.29-r1283 and those with an alignment length greater than or equal to 600 bp were extracted using SAMtools v1.13. These filtered reads were then mapped to the complete *S. nilemahensis* MAG and alignments were filtered to have a mapping quality of 30 and an alignment length greater than or equal to 600 bp. Reads from the resulting filtered alignment were determined as “shared reads” between NMV2S and *S. nilemahensis.* To determine the “total read depth” at each position of NMV2S, the original alignment of long reads mapped to NMV2S was filtered with the same quality and read length, before SAMtools v1.20 depth calculated total read coverage at each bp. The NMV2S alignment was then filtered through SAMtools v1.20 view to create a sub-alignment containing only the “shared reads” described above, but instead using their NMV2S alignment position. This alignment was again filtered for quality and read length, before being parsed through SAMtools v1.20 depth command, to generate “shared read depth”.

### Virus genome annotation

Open reading frames of viruses were predicted and annotated using Cenote-Taker3 v3.4.3^129^ in annotation mode with default parameters and without prophage pruning and wrapping. Predicted open reading frames were additionally annotated using the batch CD-Search webserver^130^ (against the CDD v3.21 database^131^ with default settings), HH-Search from HH-suite3 v3.3.0^132^ and InterProScan v5.68-100.0^71^.

Given the lack of conservation in viral sequences, structures corresponding to open reading frames were predicted using deep learning structural biology computational methods ESMfold^133^ (quay.io/nf-core/proteinfold_esmfold hash 25c14ae1b8da) and AlphaFold2 v2.3^134^ (quay.io/nf-core/proteinfold_alphafold2_split hash 4a92c9660610). Average pLDDT values were calculated for AlphaFold2 structures, and if they were below a value of 50, then the pLDDT values of the corresponding ESMfold structure was checked. If the ESMfold structure average pLDDT value was above 50 then this structure was used for subsequent analysis. Foldseek webserver^135^ was run on chosen structures with the 3Di/AA iterative algorithm (against AlphaFold/Proteome v4, AlphaFold/Swiss-Prot v4, AlphaFold/UniProt50 v4, BFVD 2023_02, CATH50 v4.3.0, PDB100 20240101 databases) with iterative search. The result with the lowest e-value was chosen if the structural alignment corresponded with a corresponding functional annotation. Annotations were only considered if e-value was less than 1e-3 and probability was above 0.9. Annotations were only accepted given previously described criteria regarding number of secondary structures in structural alignment^136^, unless the annotations were extremely consistent across databases. All sequence and structure-based annotation results were manually inspected to determine final annotation. Viral genome annotations were visualised using gbdraw^137^ and Proksee ^103^ .

Open reading frames annotated as viral structural proteins, as well as those adjacent to these structures, were further modelled as homo-multimers with varying stoichiometries using AlphaFold3^139^. Additionally, hetero-multimers were modelled based on predicted interactions between other predicted structural subunits.

AlphaFold3 was unable to confidentially model interactions between different ORF structures. Therefore PyMOL v3.1.4^140^ was used to align predicted NMV1 tail tube (ORF82) and tail sheath (ORF81) monomer structures to a resolved crystal structure of a known sheathed phage (7kh1^141^) tail complex. To compare the diameter of the siphovirus tail tube found in cryo-ET to predicted NMV2 tube structure, PyMOL v3.0.0^140^ was used to measure diameter of predicted trimeric structure of NMV2. Vacuum electrostatics for NV3 MCPs (ORF5 and ORF6), as well as resolved spindle MCP structures (7rob^36^ and 7xdiA^30^) were calculated using PyMOL v3.0.0. Additionally, *N*-link glycosylation sites on the same structures were indicated. The canonical *N*-glycosylation sequon is NX[T/S] motif where X is any amino acid except proline^36^. Structures were superimposed in PyMOL v3.0.0 and corresponding sites which all had the same NX[T/S] motif were identified and indicated. Given AlphaFold3 could not confidently model interaction between MCP ORFs (homomeric or heteromeric), NMV3 ORF5 and ORF6 monomer structures was aligned to 7xdi using PyMOL v3.1.4 align function to demonstrate how they align to the 7xdi spindle capsid structure.

### Virus population tracking

IDT PrimerQuest was used to design primers using ‘qPCR 2 Primers Interlacing Dyes’ option^142^. Primers were tested for specificity using NCBI-BLAST+ blastn^124^ against all enrichment culture contig datasets. Primers were checked for hairpins, primer homodimers and primer heterodimers using IDT’s OligoAnalyzer tool^142^. The final list of primers can be found in Supplementary Table 6.

DNA was extracted from enrichment cultures of Asgard archaea on February 26^th^ 2025. DNA was quantified using the Qubit 1× dsDNA HS kit and a Qubit Flex Fluorometer (Invitrogen). PCR reactions with EconoTaq PLUS GREEN 2X master mix and designed virus primers were used to screen the cultures for presence of the three viruses. Each 25 µL reaction consisted of 12.5 µL of master mix, 9.5 µL nuclease free water, 1 µL of forward primer (100uM), 1 µL of reverse primer (100uM) and 1 µL of template DNA. Amplification for the viral primer sets involved an initial denaturation step at 95 °C for 3 min followed by 30 cycles of denaturation at 95 °C for 30 s, annealing at 49.5 °C (NMV1) or 54 °C (NMV2 and 3) for 20 s, an elongation step at 72 °C for 15 s, and a final extension step at 72 °C for 6 min. PCR reactions were loaded into 2% agarose gels stained with SYBR Safe dye, and bands were visualized using the BioRAD Geldoc system.

### Metatranscriptomics

Quadruplicate 1 mL samples from cultures SB19.G3.3a and SB19.G3.4b were pelleted and total RNA was extracted using the New England Biolabs (NEB) Monarch Total RNA Miniprep Kit with some modifications. Briefly, the cells were lysed using 20 μL of 25 mg/mL lysozyme (MP Biomedicals) and 20 μL cold 10 mM pH 8 Tris-HCl (Sigma-Aldrich), and follow the standard protocol onwards.

RNA samples were submitted to the Ramaciotti Centre for Genomics (UNSW) for quality checks, library preparation and sequencing. The Agilent Bioanalyzer with the Agilent RNA 6000 Pico Kit were used to determine the concentration and integrity of the samples. Prior to amplification, rRNA was depleted using the Illumina Stranded Total RNA Prep with Ligation, Ribo-Zero Plus Microbiome kit. Then, the cDNA library was sequenced using paired-end on the Illumina NovaSeq 6000 system.

Once the raw read files were obtained from the Ramaciotti Centre for Genomics, firstly, the quality of the raw reads was checked using FastQC^143^ v0.11.9. Then, contaminating rRNA sequences were removed using SortMeRNA^144^ v4.3.6 with the default parameters and databases. Reads were then trimmed using the BBDuk module in BBMap (https://bbmap.org) to remove adapter sequences, reads with average Phred scores below 30, and discard any reads below 50 bp after trimming.

Using Bowtie2 v2.5.5^111^, the trimmed reads were mapped against the closed genomes of viral hosts *N. marumarumayae* and *S. nilemahensis*, NMV1–3 (including NMV2S), SNV1–4, and all other unpublished MAGs from the enrichment cultures^20^ (see above). The output, SAM files, were converted to BAM files and sorted using SAMtools v.1.15.1^112^. Then, featureCounts^145^ v2.1.1 was used to quantify the read pairs successfully aligned against NMV1–3, including NMV2S, and SNV1–4. To perform FPKM calculations, the total properly paired reads for each sample were quantified using the SAMtools v.1.15.1 flagstat command.

## Author contributions

JM, XV-C, BCM, and BPB conceptualized the project. JM maintained and monitored enrichment cultures and conducted TEM and PCR analyses. JM and XV-C performed genomic analyses including (re-)assembly of genomes, classification and annotation and undertook detailed data interpretation and visualization. MDJ and DG prepared cryo-EM samples and collected and analysed cryo-EM data. VV and DL acquired and analysed TEM data. M-CS acquired and analysed metatranscriptomic data. LH conducted specific sequence annotation analyses. BP undertook segmentation of cryo-EM data. TL and KR did the computational protein structure prediction, searches, and curation. MEP performed genomic sequencing. KAM oversaw and interpreted the computational protein structure analyses. BPB, BCF, and XV-C administered and supervised the project. BPB and DG acquired funding. JM wrote the manuscript with input and edits from all authors.

## Acknowledgements

We are grateful to Malgana language expert Kymberley Oakley, Elder Auntie Pat Oakley, and Malgana Elders for helping find a suitable name for the novel virus *Juurluvirus versutum*, and for granting permission to use the words *juurlu* from the language of the Malgana people of Gathaagudu. We acknowledge the support of the MWAC Structural Biology Facility (https://doi.org/10.26190/4KQF-M552) and UNSW Computing Cluster Katana (https://doi.org/10.26190/669x-a286) for access to computational resources. cryo-ET data were collected at the Ian Holmes Imaging Center (Bio21, University of Melbourne).

## Data availability

The raw metagenomic data and viral cMAGs will be uploaded to ENA. Cryo-ET data has been deposited in EMPIAR (deposition number TBC). Data will be made public on publication. This study did not generate any unique code.

