## Supplementary Figures 1-10 for "Viruses, Proviruses and Satellites from Asgard Archaea Enrichments Reveal Complex Microbial Interactions"

The file contains Supplementary Figures 1-10.

**Supplementary Figure 1.** Dated genealogy of the enrichment cultures used in present study.
**Supplementary Figure 2.** Montage of in silico models of NMV1 virion structural proteins.
**Supplementary Figure 3.** Montage of in silico models of all NMV2 proteins.
**Supplementary Figure 4.** Montage of in silico models of all NMV2S proteins.
**Supplementary Figure 5.** Montage of in silico models of all NMV3 proteins.
**Supplementary Figure 6.** PCR results of NMV1–3 in enrichment cultures.
**Supplementary Figure 7.** Montage of in silico models of SNV1 virion structural proteins.
**Supplementary Figure 8.** Montage of in silico models of SNV2 virion structural proteins.
**Supplementary Figure 9.** Montage of in silico models of SNV3 virion structural proteins.
**Supplementary Figure 10.** Montage of in silico models of all SNV4 proteins.

**Other Supplementary Materials for this manuscript include the following:**

Supplementary Tables 1-10; Supplementary Video 1

**Supplementary Table 1.** Summary of *Nerearchaeum marumarumayae* and *Stromatodesulfovibrio nilemahensis* (pro)viruses
**Supplementary Table 2.** Nerearchaeum marumarumayae  CRISPR Spacer Blast Matches
**Supplementary Table 3.** Hi-C host-virus linkages
**Supplementary Table 4.** Final annotations of NMVs and SNVs
**Supplementary Table 5.** FPKM values per NMV and SNV genes
**Supplementary Table 6.** PCR primer sequences
**Supplementary Table 7.** Correlation values between NMVs and host
**Supplementary Table 8.** vConTACT3 final taxonomic assignments
**Supplementary Table 9.** Previously described Non-Refseq Genomes included in vContact3 analysis
**Supplementary Table 10.** Scaffolds from IMG/V database included in analysis

**Supplementary Video 1.**  3D tomogram reconstruction and segmentation of *Nerearchaeum marumarumayae* and associated viruses

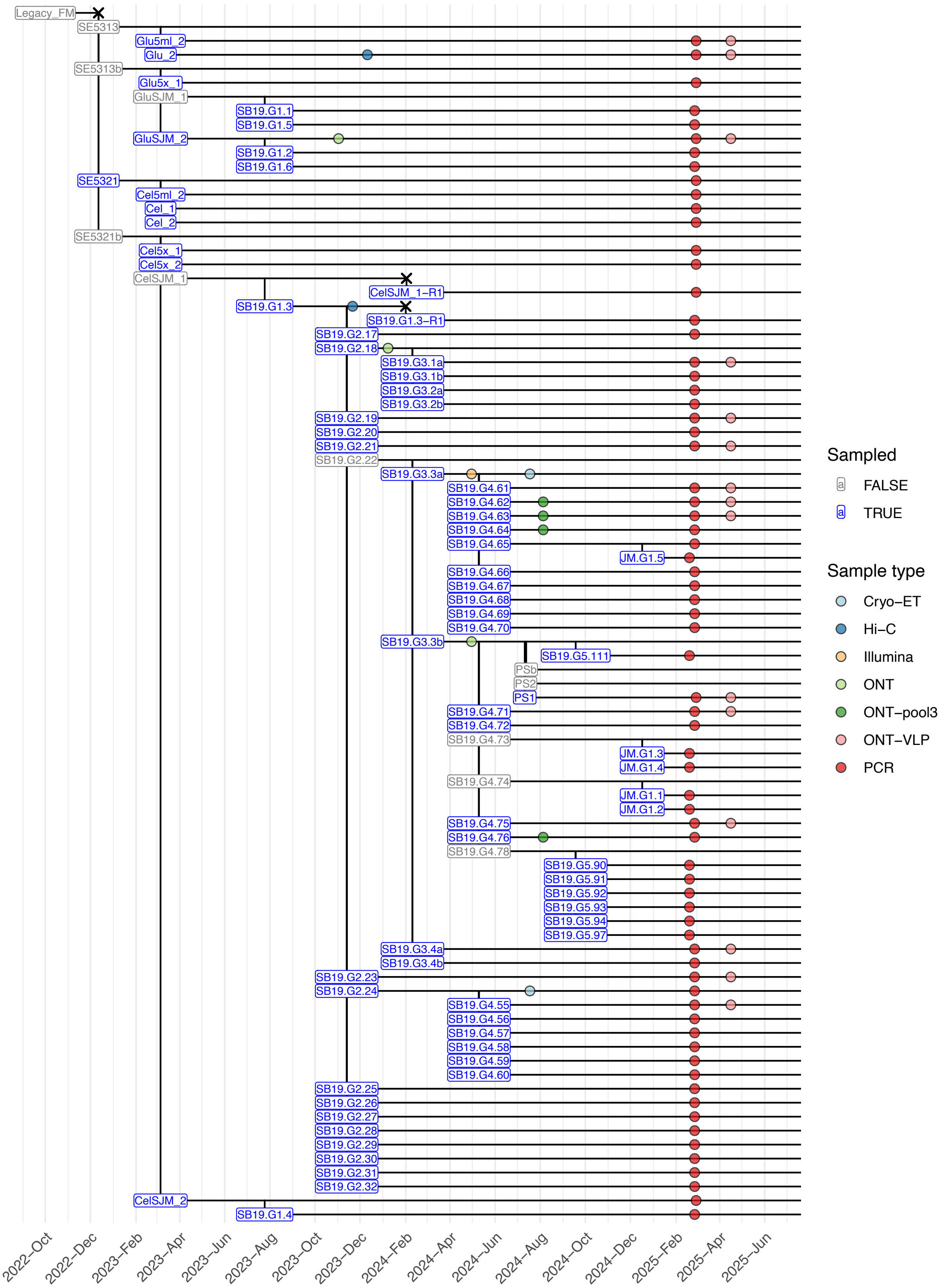


**Supplementary Figure 1. Dated genealogy of the enrichment cultures used in present study.** Names of sampled cultures appear in blue. Coloured circles indicate the usage of the samples. All names and sampling points are centred on the collection/inoculation date. Crosses mark culture end dates. See Methods for description of sample types. Dates for cultures prior to 2023 are approximate.

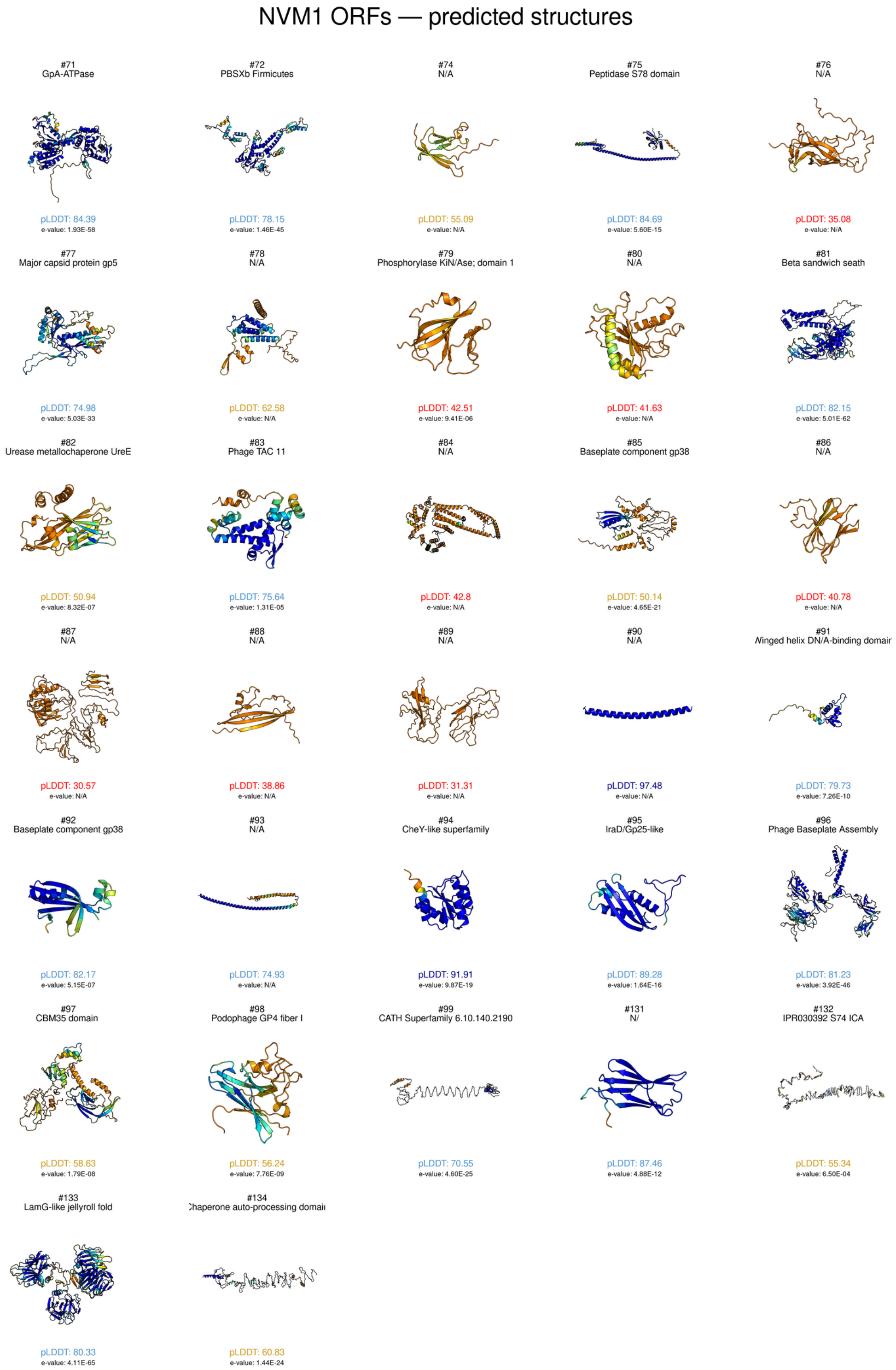


**Supplementary Figure 2. Montage of in silico models of NMV1 virion structural proteins.** In silico AlphaFold structures predicted from protein sequences of predicted virion structural proteins of NMV1 in this study. The residue position confidence scores are reported, coloured by the pLDDT (the predicted local distance difference test) across the whole protein. Dark blue scores are considered to have good global backbone and residue positioning, light blue indicates confidence in the global fold but not detailed positioning, yellow indicates low confidence across the prediction, and red values demonstrate uninterpretable predictions or disordered regions.

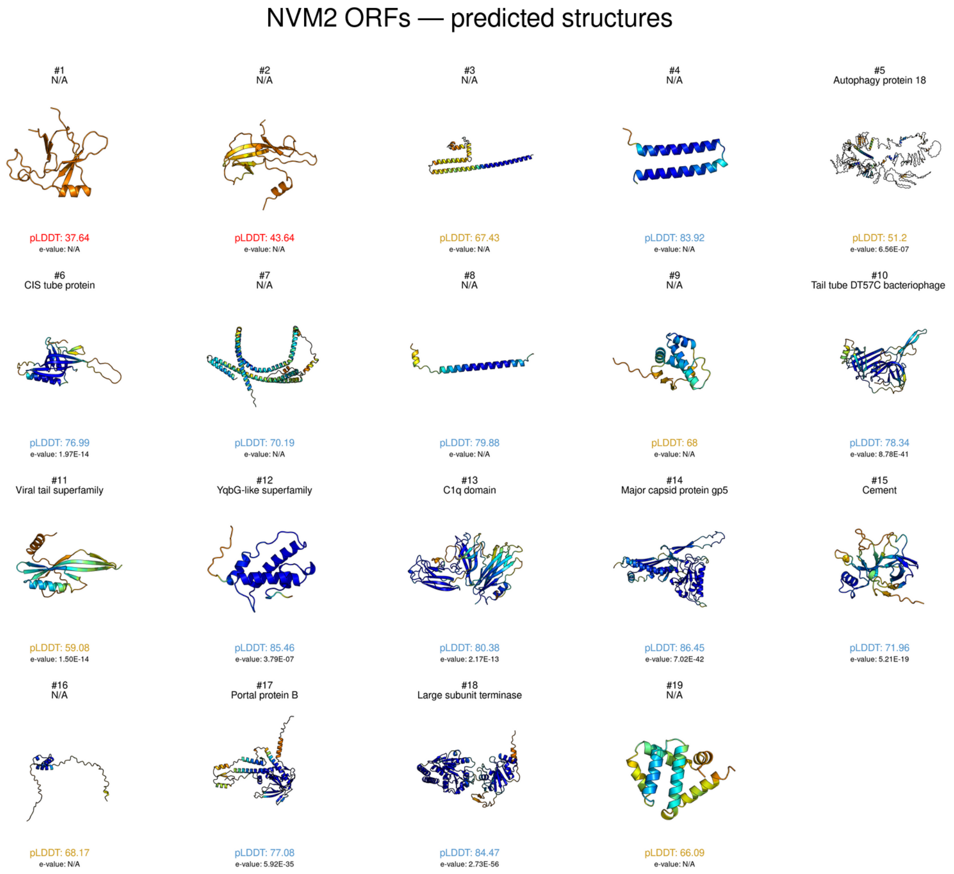


**Supplementary Figure 3. Montage of in silico models of all NMV2 proteins.** In silico AlphaFold structures predicted from protein sequences of all proteins of NMV2 in this study. The residue position confidence scores are reported, coloured by the pLDDT (the predicted local distance difference test) across the whole protein. Dark blue scores are considered to have good global backbone and residue positioning, light blue indicates confidence in the global fold but not detailed positioning, yellow indicates low confidence across the prediction, and red values demonstrate uninterpretable predictions or disordered regions.

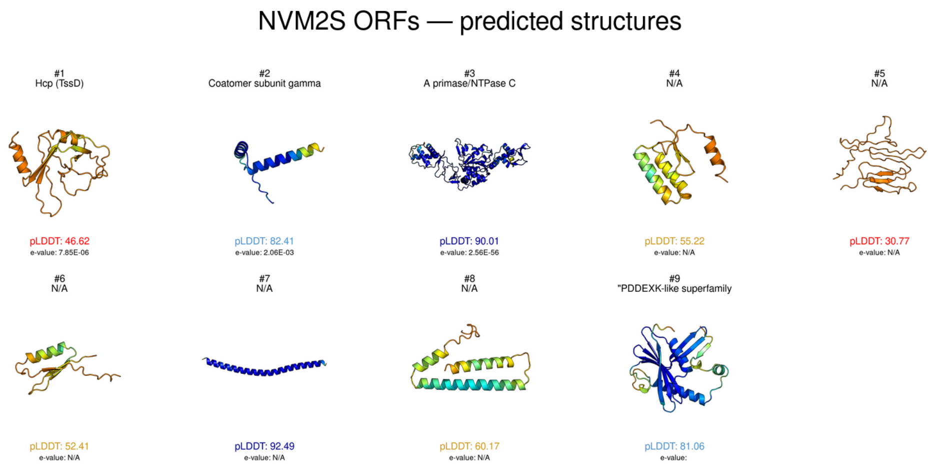


**Supplementary Figure 4. Montage of in silico models of all NMV2S proteins.** In silico AlphaFold structures predicted from protein sequences of all proteins of NMV2S and NMV3 in this study. The residue position confidence scores are reported, coloured by the pLDDT (the predicted local distance difference test) across the whole protein. Dark blue scores are considered to have good global backbone and residue positioning, light blue indicates confidence in the global fold but not detailed positioning, yellow indicates low confidence across the prediction, and red values demonstrate uninterpretable predictions or disordered regions.

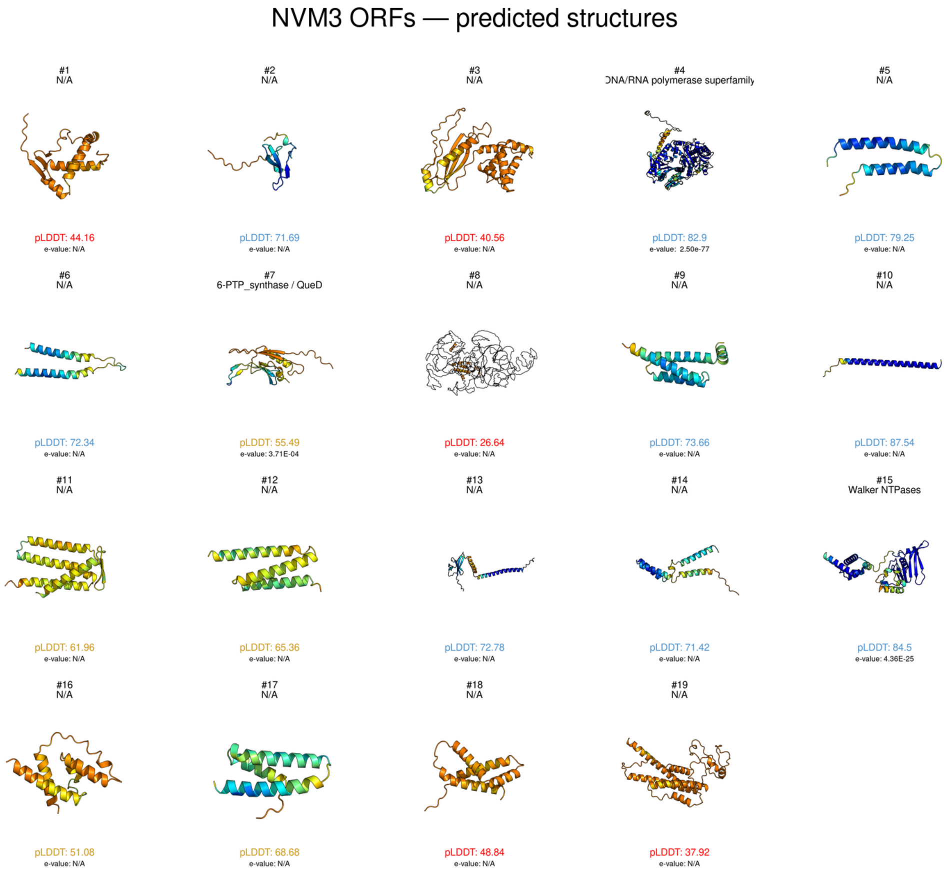


**Supplementary Figure 5. Montage of in silico models of all NMV3 proteins.** In silico AlphaFold structures predicted from protein sequences of all proteins of NMV2S and NMV3 in this study. The residue position confidence scores are reported, coloured by the pLDDT (the predicted local distance difference test) across the whole protein. Dark blue scores are considered to have good global backbone and residue positioning, light blue indicates confidence in the global fold but not detailed positioning, yellow indicates low confidence across the prediction, and red values demonstrate uninterpretable predictions or disordered regions.

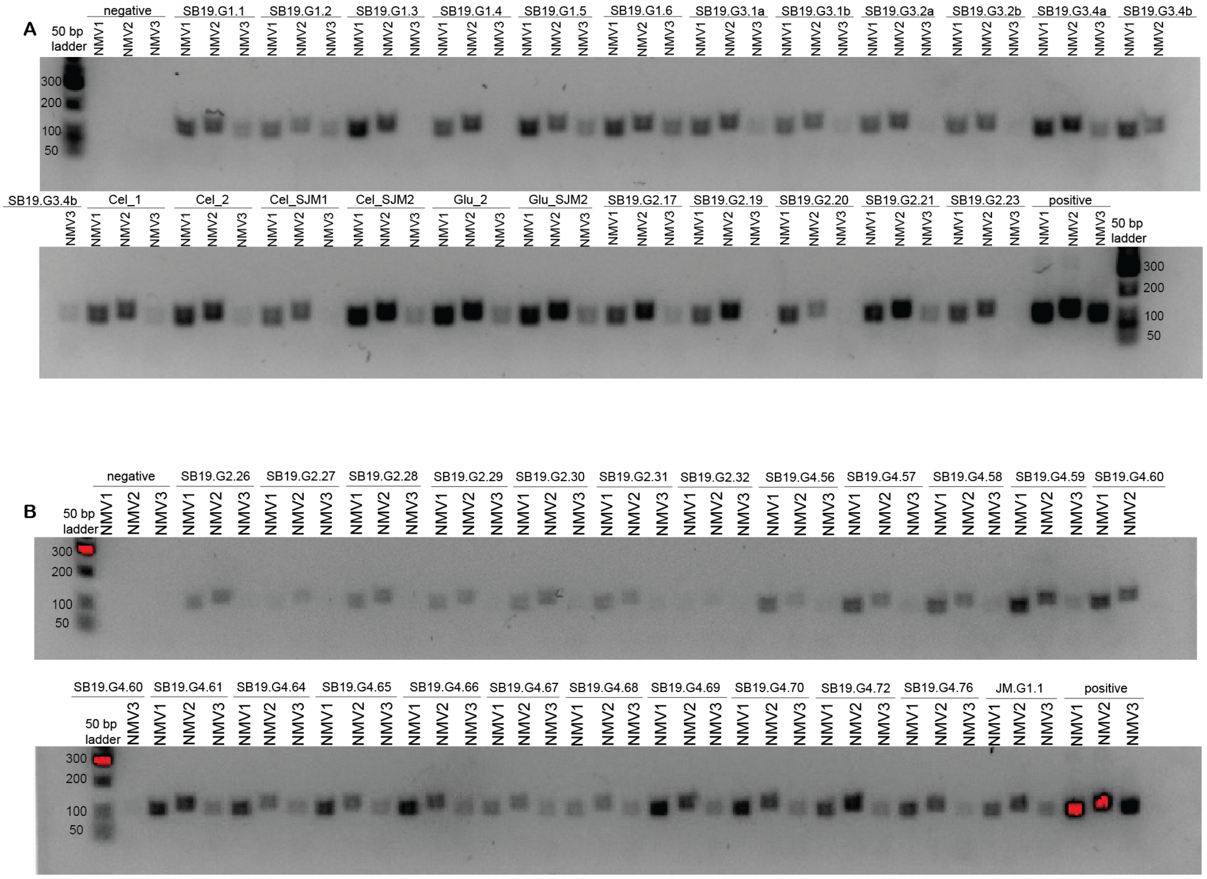


**Supplementary Figure 6. PCR results of NMV1–3 in enrichment cultures.** Agarose gel electrophoresis of PCR products generated using NMV-specific primers for NMV1–3 as specified in Supplementary Table 6. Three different gels (A and B) of the same PCR reaction are shown. Lanes are labelled as the enrichment culture DNA originated from and the virus targeted.  Expected size of products is detailed in Supplementary Table 6 with bands of correct size shown in positive controls, and an absence of bands shown in negative controls.

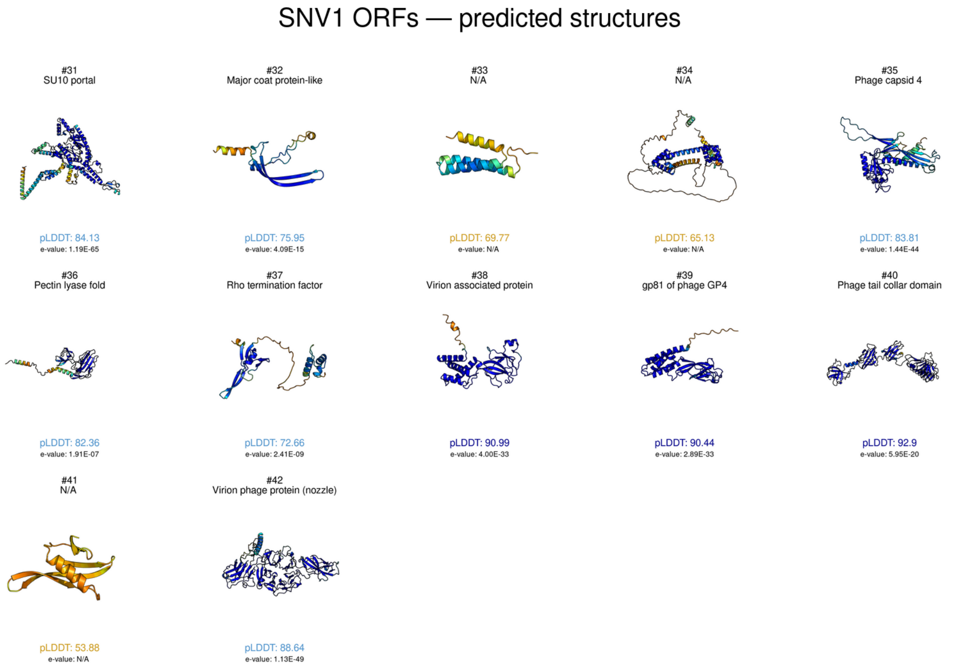


**Supplementary Figure 7. Montage of in silico models of SNV1 virion structural proteins.** In silico AlphaFold structures predicted from protein sequences of predicted virion structural proteins of SNV1 in this study. The residue position confidence scores are reported, coloured by the pLDDT (the predicted local distance difference test) across the whole protein. Dark blue scores are considered to have good global backbone and residue positioning, light blue indicates confidence in the global fold but not detailed positioning, yellow indicates low confidence across the prediction, and red values demonstrate uninterpretable predictions or disordered regions.

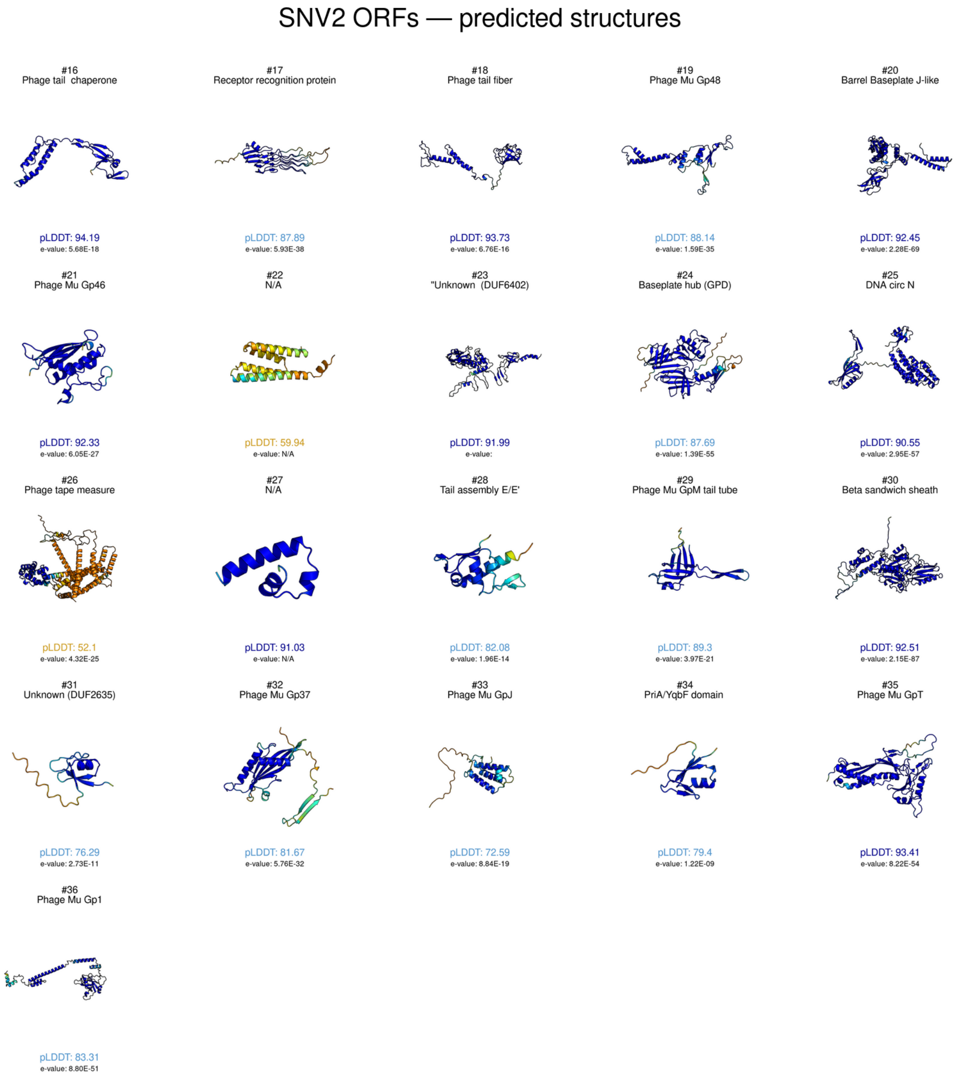


**Supplementary Figure 8. Montage of in silico models of SNV2 Vir virion structural proteins.** In silico AlphaFold structures predicted from protein sequences of predicted virion structural proteins of SNV2 in this study. The residue position confidence scores are reported, coloured by the pLDDT (the predicted local distance difference test) across the whole protein. Dark blue scores are considered to have good global backbone and residue positioning, light blue indicates confidence in the global fold but not detailed positioning, yellow indicates low confidence across the prediction, and red values demonstrate uninterpretable predictions or disordered regions.

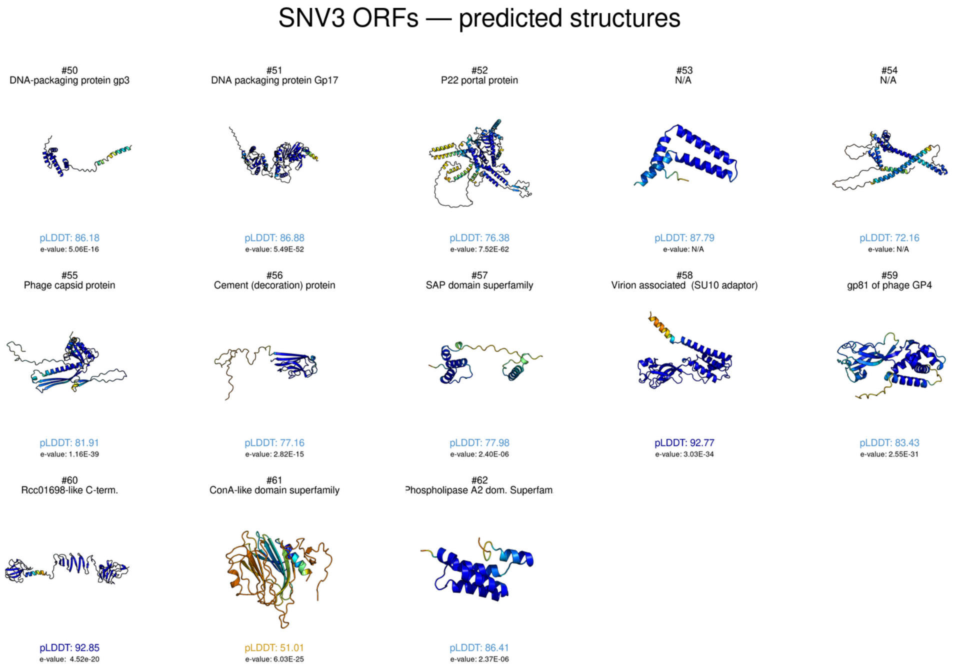


**Supplementary Figure 9. Montage of in silico models of SNV3 virion structural proteins.** In silico AlphaFold structures predicted from protein sequences of predicted virion structural proteins of SNV3 in this study. The residue position confidence scores are reported, coloured by the pLDDT (the predicted local distance difference test) across the whole protein. Dark blue scores are considered to have good global backbone and residue positioning, light blue indicates confidence in the global fold but not detailed positioning, yellow indicates low confidence across the prediction, and red values demonstrate uninterpretable predictions or disordered regions.

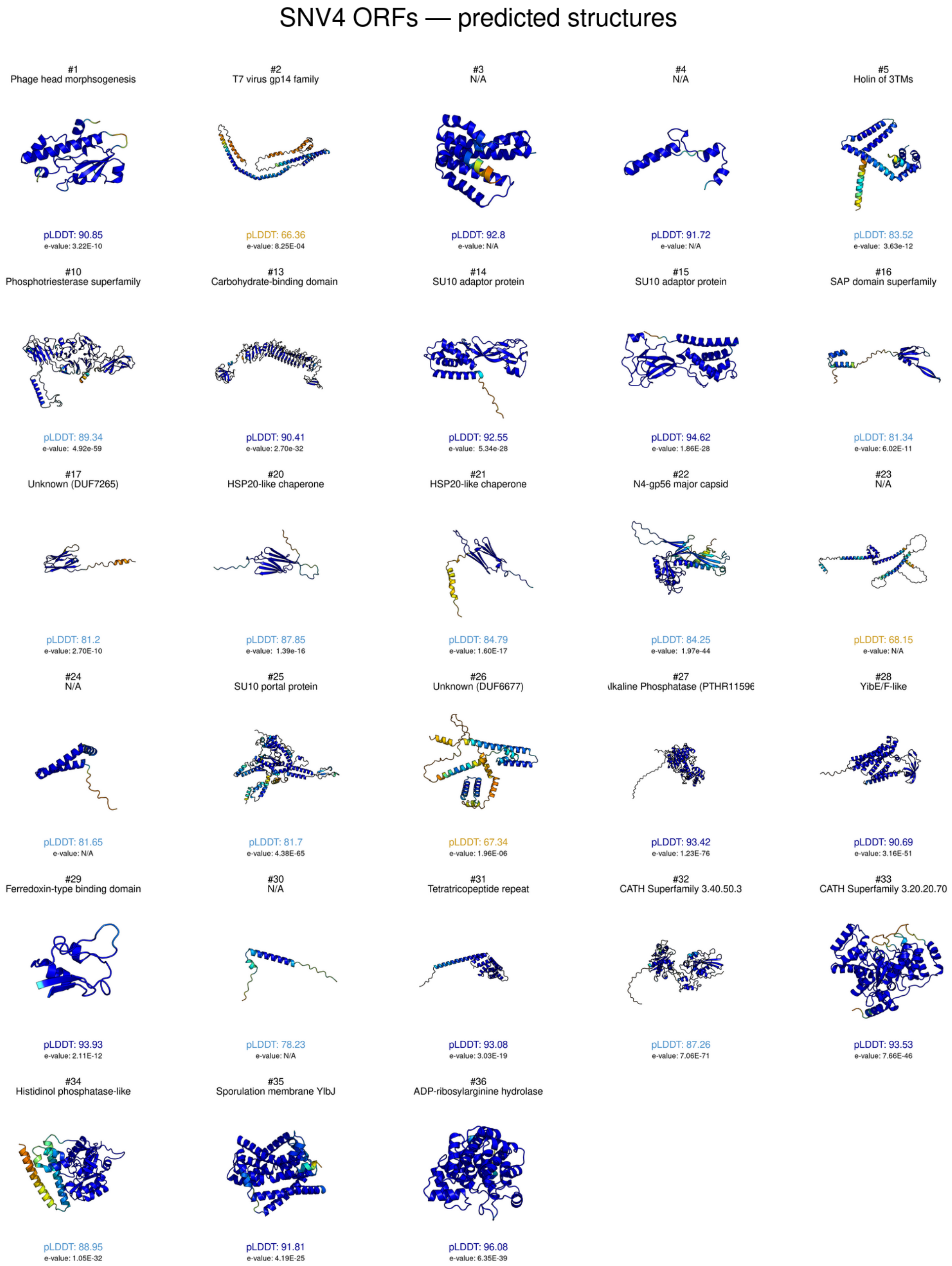


**Supplementary Figure 10. Montage of in silico models of all SNV4 proteins.** In silico AlphaFold structures predicted from protein sequences of predicted proteins of SNV4 in this study. The residue position confidence scores are reported, coloured by the pLDDT (the predicted local distance difference test) across the whole protein. Dark blue scores are considered to have good global backbone and residue positioning, light blue indicates confidence in the global fold but not detailed positioning, yellow indicates low confidence across the prediction, and red values demonstrate uninterpretable predictions or disordered regions.
