## Extended Data Figures 1-3 for "Viruses, Proviruses and Satellites from Asgard Archaea Enrichments Reveal Complex Microbial Interactions"

**Extended Data (Meltzer et al)**

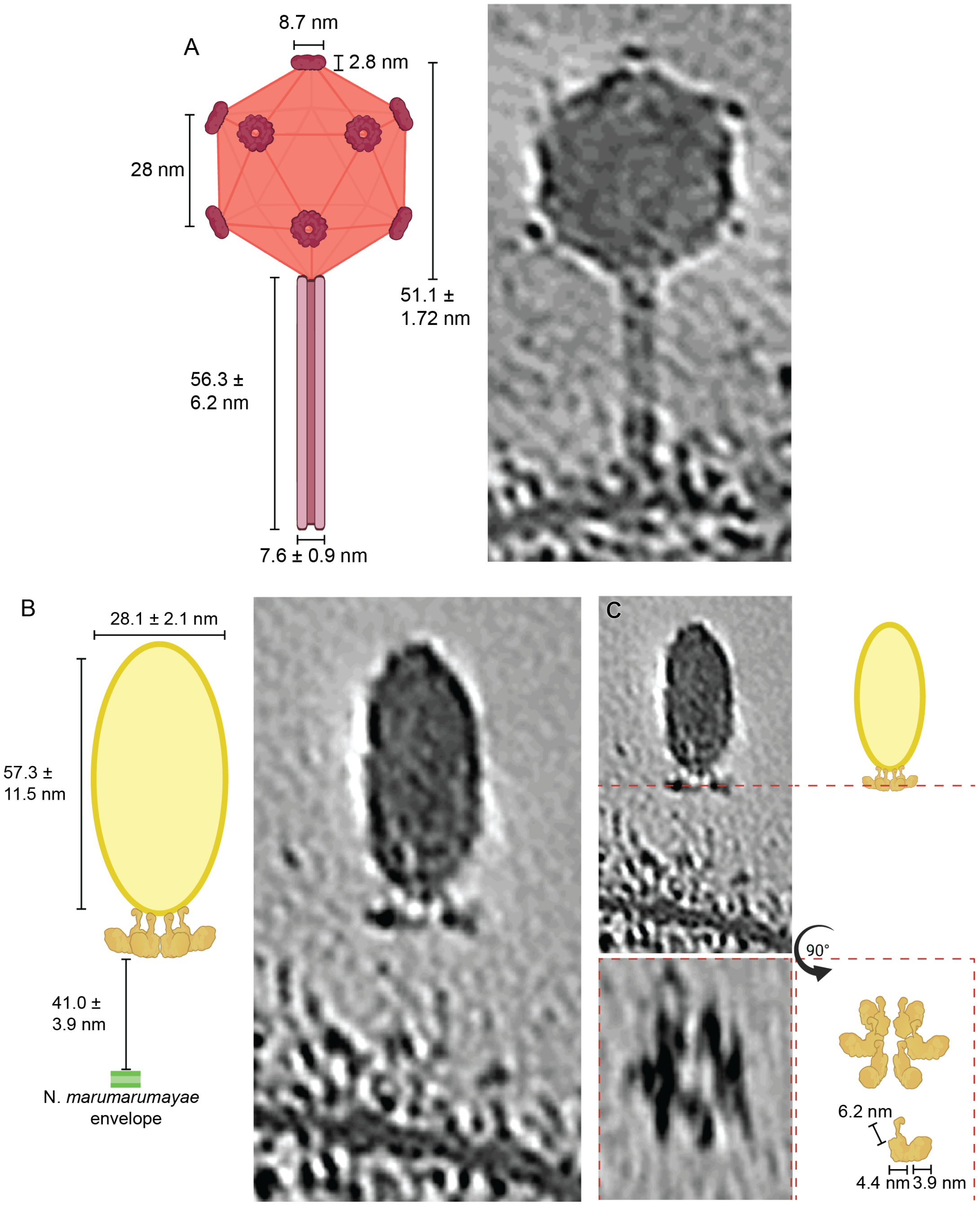


**Extended Data Figure 1. Measurements and details of predicted virus morphotypes in cryo-ET**. Cryo-ET image of siphovirus **(A)** and spindle virus **(B)** morphotype with corresponding cartoon representation of predicted morphology with measurements **(C)**. Wedge-like structure observed in spindle virus morphotype is shown in more detail across a 90° rotation of the tomogram (C).

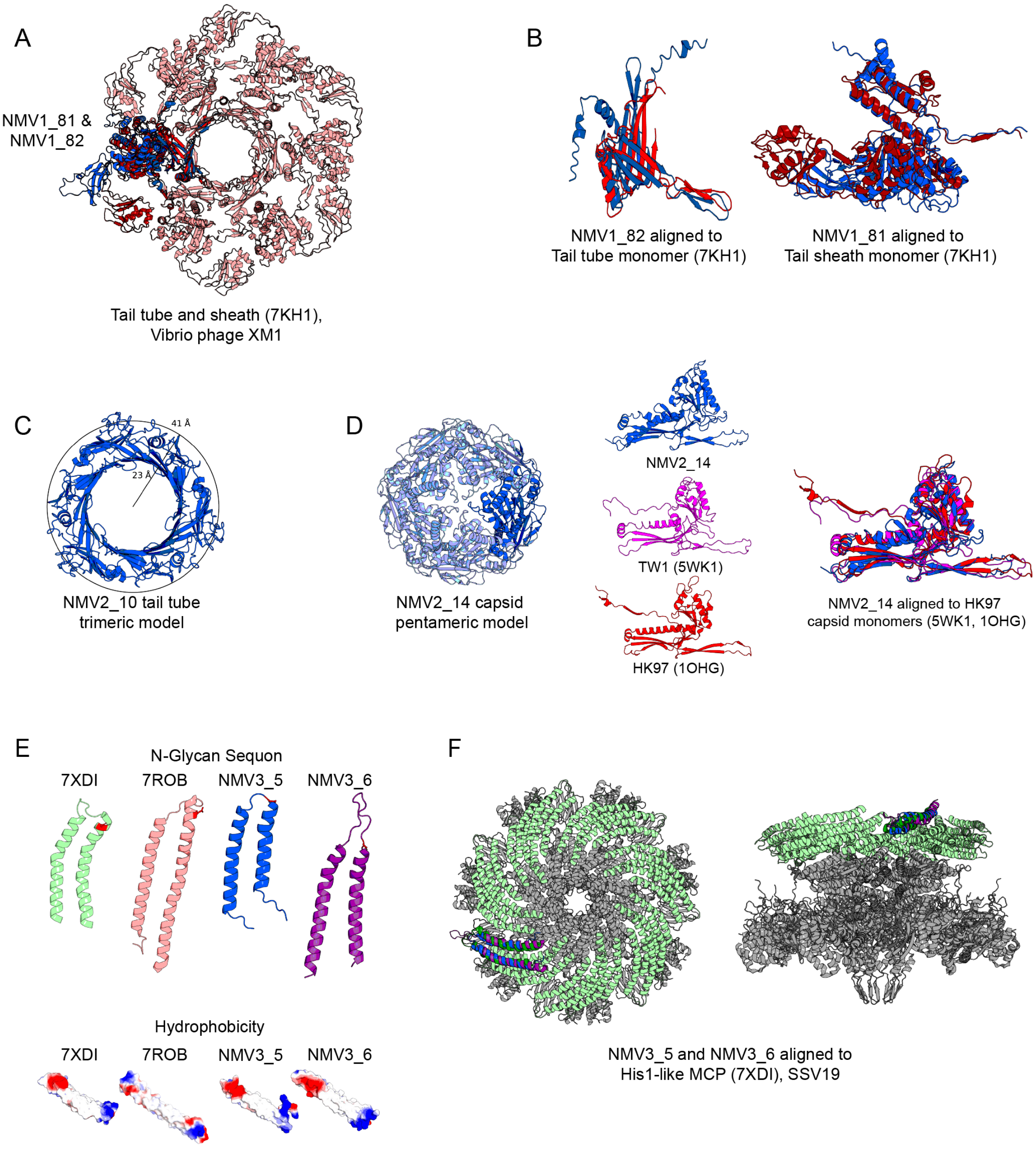


**Extended Data Figure 2. Detailed analysis and comparisons of modelled hallmarks of predicted NMV morphotypes. (A)** Modelled structures of NMV1 tail tube and sheath monomers (coloured blue) aligned to previously resolved crystalline structures (7KH1) of a sheathed tail (complex in salmon and monomer in alignment coloured red). **(B)** NMV1 tail tube and sheath monomers aligned to the monomers of tube and sheath from 7KH1. **(C)** Trimeric model of NMV2 tail protein with dimensions of inner and outer radius. Pentameric model of NMV2 capsid with a monomer highlighted. **(D)** Monomer is compared to previously described resolved crystalline structures of HK97 fold capsid monomers. **(E)** Resolved known spindle capsid monomer structures (7XDI, 7ROB) beside modelled NMV3 capsid monomer structures with *N*-glycan sequon sites highlighted in red, with capsid structures are coloured based on vacuum electrostatics (bottom). **(F)** NMV3 ORF 5 monomer (blue) and NMV3 ORF 6 monomer (pink) are aligned to the SSV19 MCP structure (7XDI) (green), with tail structure shown in grey.

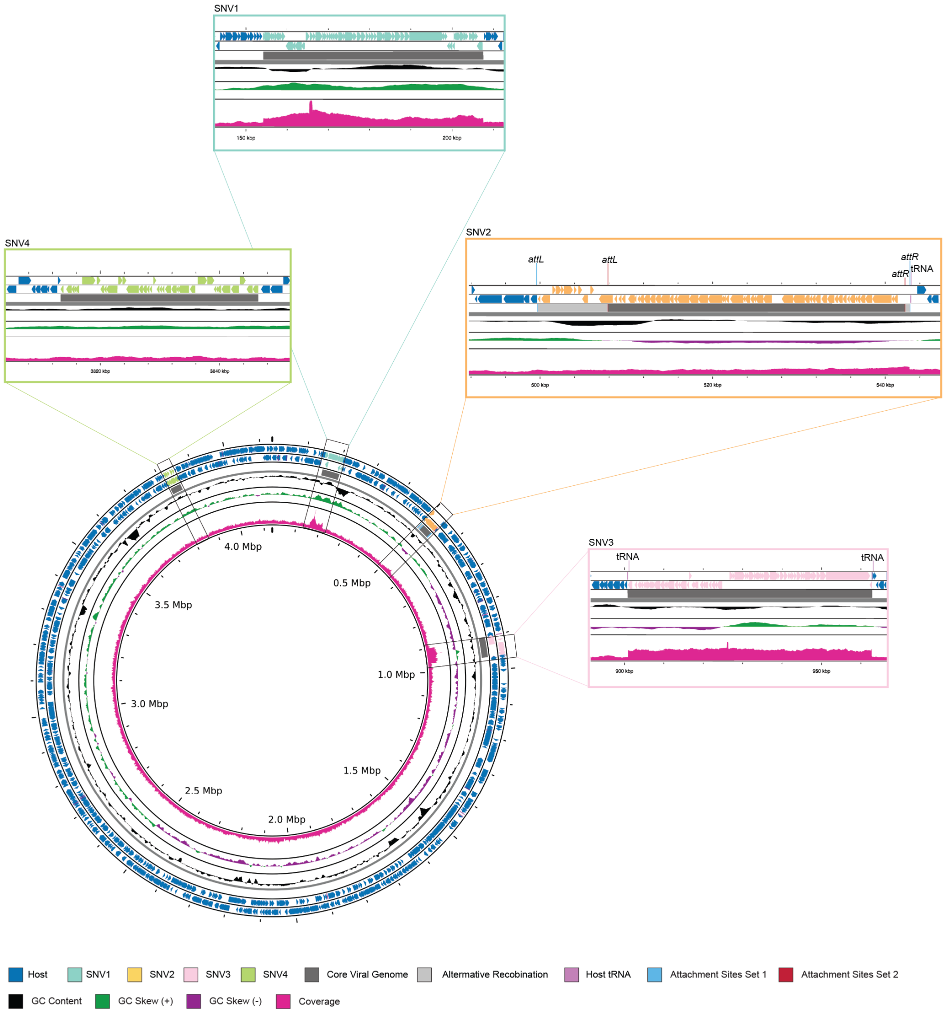


**Extended Data Figure 3. Predicted viral Boundaries of SNVs.** Shown here are the marked changes in genome coverage of SNV1 and SNV3 as well as the predicted attachment and alternative recombination sites for SNV2.
